# Time cells differentially populate trace and post-trace epochs but do not remap for different trace intervals

**DOI:** 10.64898/2026.02.26.708289

**Authors:** Hrishikesh Nambisan, Upinder S. Bhalla

## Abstract

Hippocampal time cells are a neural substrate for linking discontiguous events in time. While they have been reported both for working memory tasks (∼15 s) and trace-eyeblink conditioning (TEC, ∼300 ms), the latter is ∼50 fold faster and requires pre-emptive and precisely timed responses. We therefore asked if novel features of time-cell rescaling, post-stimulus activity, and time-cell extinction might arise in TEC. We used 2-photon calcium-sensor imaging from mouse hippocampal CA1 pyramidal neurons during TEC tasks with varying intervals, and specifically included a protocol in which short blocks of 250 and 550ms trials were alternated to eliminate learning-dependent rescaling. We find that conditioned stimulus (CS) triggered traces extend for >5 seconds after the unconditioned stimulus (US), and the trials can be subdivided into epochs with distinct proportions of time cells and time-cell peak widths. Instead of remapping, the time-cell sequence is the same between 250 and 550ms intervals. CS-triggered time cells are present before trace learning, but are nearly lost after behavioural extinction, suggesting that their removal is an active process. We propose that in TEC, salient stimuli initiate time-cell sequences which are independent of subsequent timing or even absence of US, but can be gated off during extinction.

## Introduction

The hippocampus forms representations of linear and multidimensional context spaces, most notably for space, but including time and sensori-motor axes when the task demands (Hasselmo & Stern, 2015; Howard et al., 2014; Howard & Eichenbaum, 2013, 2015). A common feature of each of these representations is that one can decode the relevant context from the activity of a population of hippocampal neurons. For example, by monitoring enough CA1 neurons in a mouse, one can ascertain where the mouse is currently situated in an arena (Guger et al., 2011). Furthermore, when neural representations of space or time are experimentally perturbed, the animal exhibits altered behaviour consistent with the interpretation that the animal’s sense of space and time is mechanistically derived from this contextual structure of population activity (Eichenbaum, 2014, 2017).

### Dynamic allocation of cells for representations of space and time

The representations of different contexts utilize overlapping sets of cells, suggesting that the hippocampus deploys its cellular resources dynamically. Specifically, the same cell may function as a time cell in a time-dependent learning task, and as a place cell when the animal is exposed to a spatial arena (Kraus et al., 2013). Several key predictions emerge from this dynamic allocation of resources: that space and time representations should be learnt, that they should remap when the context is entirely different, that they should change (stretch or morph) when the spatial arena is slightly altered or task duration is extended, and that they should be lost when the task loses relevance. Below we compare observations for place and time cells in each of these four domains.

### Place and time cell learning

Place cells are indeed learnt, and this learning takes place on remarkably few exposures (Hill, 1978; Wilson & McNaughton, 1993). Using intracellular recordings from animals learning a new arena, Bittner et al found that several places elicit subthreshold CA1 PN depolarization, suggesting the existence of multiple stimulus-driven latent place fields (Bittner et al., 2015). When one such depolarization crossed threshold into spiking, either spontaneously or driven by the experimenter, there was rapid BTSP potentiation and within a few trials the place field of the cell stabilized at the selected location (Bittner et al., 2017). For time cells, work from Buzaki and Eichenbaum’s groups (MacDonald et al., 2011; Pastalkova et al., 2008) and (Modi et al., 2014)) report creation of new time cells during learning, though the learning relevance of time cells has been disputed (Sabariego et al., 2019; Salz et al., 2016). Modi et al.(Modi et al., 2014) reported that before trace-eyeblink conditioning, the activity of prospective time cells was dispersed, but converged to a tight time-cell peak upon task acquisition. Using weaker conditioned stimuli in a similar trace-eyeblink task, we have also seen emergence of stimulus triggered time-cells prior to task learning (Bhattacharjee et al., 2024). This has parallels with proposals of spontaneous activity sequences in the hippocampus, which can be linked to time-relevant events (Dragoi & Buzsáki, 2006; Dragoi & Tonegawa, 2011; Villette et al., 2015)

### Remapping

Place cells remap rapidly in different arenas (Muller & Kubie, 1987) on a time-scale similar to that of the original learning (Bostock et al., 1991; Wilson & McNaughton, 1993). There are further context-specific nuances. For example, there are ‘splitter’ cells which map exactly the same stretch of linear track differently depending on context, such as whether the animal has to turn left or right at the end of the linear stretch (Frank et al., 2000; Wood et al., 2000). Eichenbaum has shown that task salience, such as reward status, also alters place cell representations (Komorowski et al., 2009; Lee et al., 2006). For rescaling of linear tracks there is a range of outcomes: border cells at the start and end stay mostly fixed, but many of the intermediate cells stretch (O’Keefe & Burgess, 1996). Additionally some cells may change their field (Muller & Kubie, 1987; O’Keefe & Burgess, 1996). Relatively few studies have been done on representation changes in the time-cell domain. Time-cells undergo a range of effects, from constancy, shifting, to new cell recruitment, when the interval in an object-odor mapping task is altered (Cao et al., 2022; MacDonald et al., 2011) or when the nature of the task changes (MacDonald et al., 2013). Remapping of time cells has also been seen in human MTL when the task switches between search and retrieval (Umbach et al., 2020). In animals, distinct odor memories form distinct time cell sequences (MacDonald et al., 2013), and likewise sound and light stimulus modalities result in distinct but partially overlapping populations of time cells (Bhattacharjee et al., 2024). To our knowledge no studies have examined time-cell rescaling for the much shorter intervals of hippocampal trace conditioning.

### Loss and extinction of representations

Extinction of place cell representation, in the classical behavioural sense, is not supported in the literature. Some features of loss of representation take the form of place cell drift over repeated exposure (Mankin et al., 2012; Ziv et al., 2013). In contrast, many cells retain their place location over a period of weeks to months (Lt & Pj, 1990). Given that place cell maps form in the absence of overt behavioural reinforcement, the lack of extinction may not be surprising, though it is remarkable how the hippocampus maintains long, stable, and numerous representations of place (Lever et al., 2002; Lt & Pj, 1990; Ziv et al., 2013). Time-cells, in contrast, are typically bracketed by a stimulus and a reinforcement. Nevertheless, extinction of time-cells representation may be similar to that in place cells, in that it is a combination of representational drift (Bhattacharjee et al., 2024; Mau et al., 2018) and remapping (Masuda et al., 2020).

In the current study we address four key gaps in the understanding of time-cells, with particular reference to TEC: First, are rescaling properties different between multi-second (working memory) tasks compared to sub-second (trace eyeblink conditioning) behavior? This addresses the mechanisms of time-cell formation and remapping over very different time domains. Second, are time-cell sequences in TEC contingent on learning an association, and once learnt, are they dependent on the timing of the US? This brings the TEC perspective to an ongoing debate in the time-cell field. Third, how does the time-cell sequence evolve over times much longer than the trace period? This refers to the relationship between event triggers, intrinsic brain sequences and their evolution in time. Finally, do post-extinction time cells fall back to a pre-learning state? This is particularly applicable in our study because we observe time cells before learning.

## Results

We trained 7 mice on a staged trace-eyeblink conditioning task with increasing trace intervals from 250 to 550 ms (Figure 1A-E). We then challenged them with an interleaved task having alternating blocks of five trials each of 250 and 550 ms. Finally, we removed the air-puff reinforcement to elicit extinction. Throughout this process we recorded activity from the dorsal hippocampal CA1 pyramidal neurons of 4 of the trained mice using 2-photon imaging of Ca responses reported by GCAMP-6f (Figure 1F, G). We compared network activity as well as time-cell activity across each of these stages, and contrasted them between the long and short trace periods for interleaved trials. We also monitored treadmill activity and did not find any correlation with neuronal activity (supplementary Figure 1).

**Figure 1:**
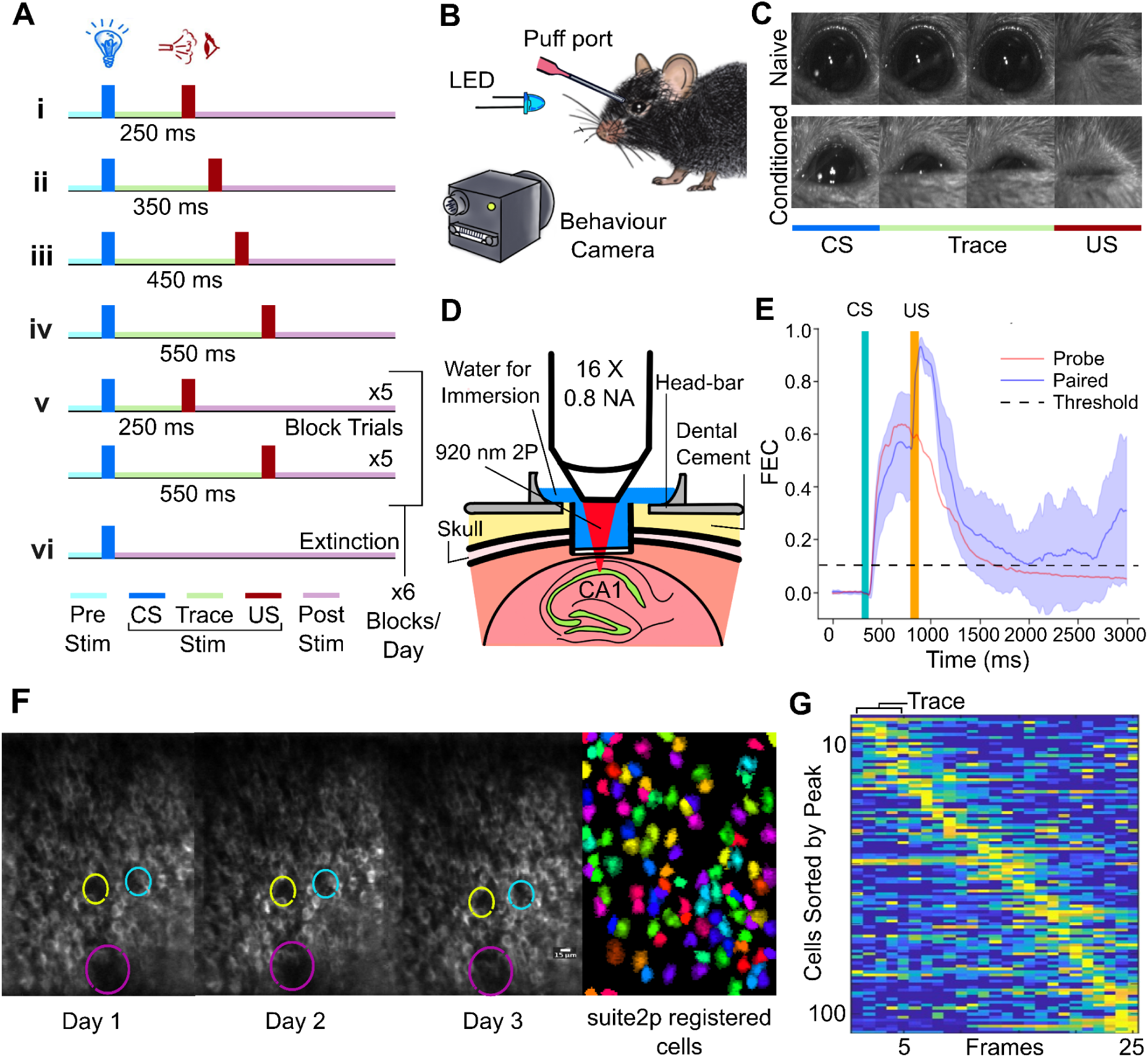
Overview of setup and baseline data. **A:** Staged structure of trace eyeblink conditioning task. The CS was a 50 ms flash from a blue LED and the US a 50 ms air puff to the eye. Stages i to vi represent 250ms trace, 350ms, 450ms, 550ms, interleaved, and extinction trials. **B:** Schematic of stimulus delivery and monitoring. **C:** Video frames from the behaviour camera recorded at 200fps during the task. Above row: before learning. Below: after learning. Note that the animal starts to shut its eye before the arrival of the air puff once it has learned the task. **D:** Cartoon schematic of optical imaging. A cannula with a flat glass coverslip below was placed just above the left or right hippocampus after removing a small amount of somatosensory cortex. **E:** Example Fraction Eye Closure (FEC) profile from a single session of an animal that has learnt the association, as indicated by the rise in FEC value during the trace period for both the paired and probe trials. **F:** Example 2P imaged frames for three consecutive days with landmarks (indicated in coloured circle) used to track the same ROI, and after cells registered using Suite2p. **G:** Heatmap of cell activity over 25 frames sorted by peak activity, without selecting for time cells, 2P frame rate = 12.8 Hz. The stimulus is delivered on the first frame.

### Mice learn the TEC task for all intervals and tune their eyeblink times to task demands

Seven animals successfully learnt the task (Figure 2 A). We staged the trace interval starting at 250 ms since that has been reported to be the easiest for mice to learn (Siegel et al., 2015; Tseng et al., 2004). This first stage of learning took 1 to 7 days to cross the criterion of 60% of trials eliciting >10% eyeblink, following which the learning transferred immediately to longer intervals (Figure 2A). As expected, eyeblinks were progressively lost during the extinction phase. The peak of eyeblink closure closely tracked the trace interval (Figure 2B) but interestingly blink latency (time before 10% eye closure) did not change much (Figure 2C). We separated these analyses for the 250ms and 550ms intervals of the interleaved trials (Figure 2D-F). While there was a slight improvement in performance for the 550ms compared to 250ms trials (Wilcoxon test, p=0.0017, Figure 2D), the blink peak was strikingly different (Mann-Whitney U, p<1e-41, Figure 2E). This indicates that the animals were reliably tuning their eyeblinks to the trace intervals. To test if the animals began their eyeblinks according to the trace interval, we plotted the blink latency as a function of trial number within the 5-trial blocks of long and short traces. There was a small difference in blink latency for the 250 and 550ms trials (Mann-Whitney U, p=0.037, Figure 2F).

**Fig. 2:**
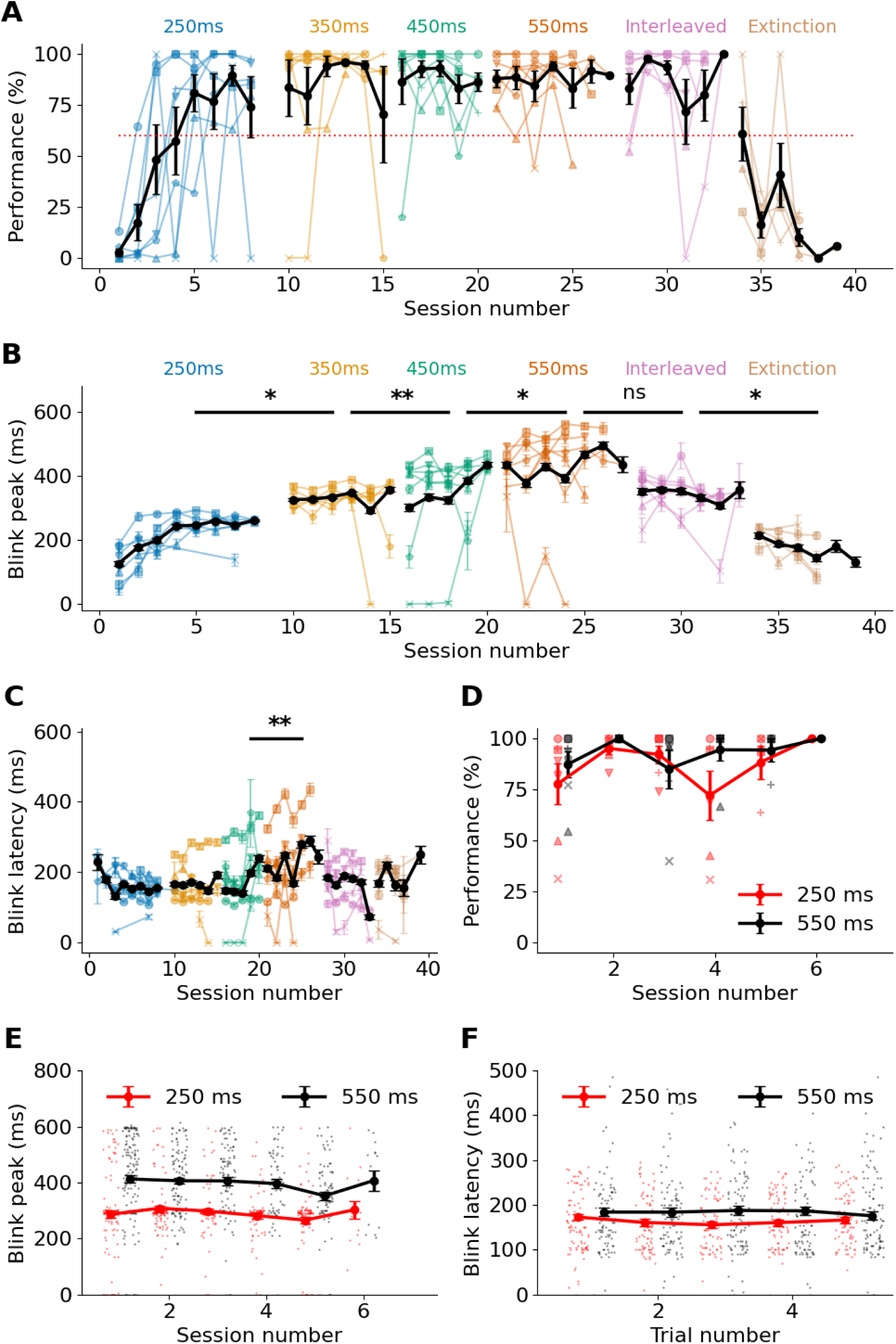
Behaviour over the different phases of training. **A:** Animal performance as a function of session number, reported as % of trials in which eyeblink occurs before US. The different colors represent different session types. Each animal is represented by a different trace. Black is the average performance. Criterion = 60% is indicated by a red dashed line. **B:** Peak blink time this rises with the trace interval, then declines for interleaved and extinction blocks (paired t-test between successive session types: p=0.049, p=0.01, p=0.01, p=0.15, and p=0.025) **C:** Latency for eyeblink, defined as the first frame in a trial in which eye closure exceeds 10%. This is mostly unchanged across sessions (paired t-test between successive session types: p = 0.78, p=0.05, p=0.008, p=0.37, p=0.9). **D:** Performance in interleaved trials. Performance is slightly better in 550ms trials (Wilcoxon test, p = 0.0017). **E:** Blink peaks are well separated for 250 and 550 ms trace intervals for all sessions where there were interleaved trials (Mann-Whitney U 62851, p<1e-41). **F:** Blink latency plotted against trial number for interleaved sessions. It is slightly but significantly separated (Mann-Whitney U 88200, p=0.036).

### Activity and time-cells increase post-stimulus, but proportions do not change over learning

We characterized network activity as mean numbers of calcium events defined as activity reaching 2 SD above baseline calculated during pre-stimulus period, per cell, per frame. We analyzed these 2P recordings to obtain reliable time cells as per the criteria from Modi et al 2014 (Figure 3A, Ananthamurthy & Bhall (2023)), using a fast C++ implementation of the algorithm (Ananthamurthy and Bhalla, 2023). We defined five epochs for each trial: Pre-CS (Epoch 0), CS to CS+1 (Epoch 1), CS+1 to CS+3 (Epoch2), CS+3 to CS+6 (Epoch 3) and >CS+6 (Epoch 4).

**Fig. 3.**
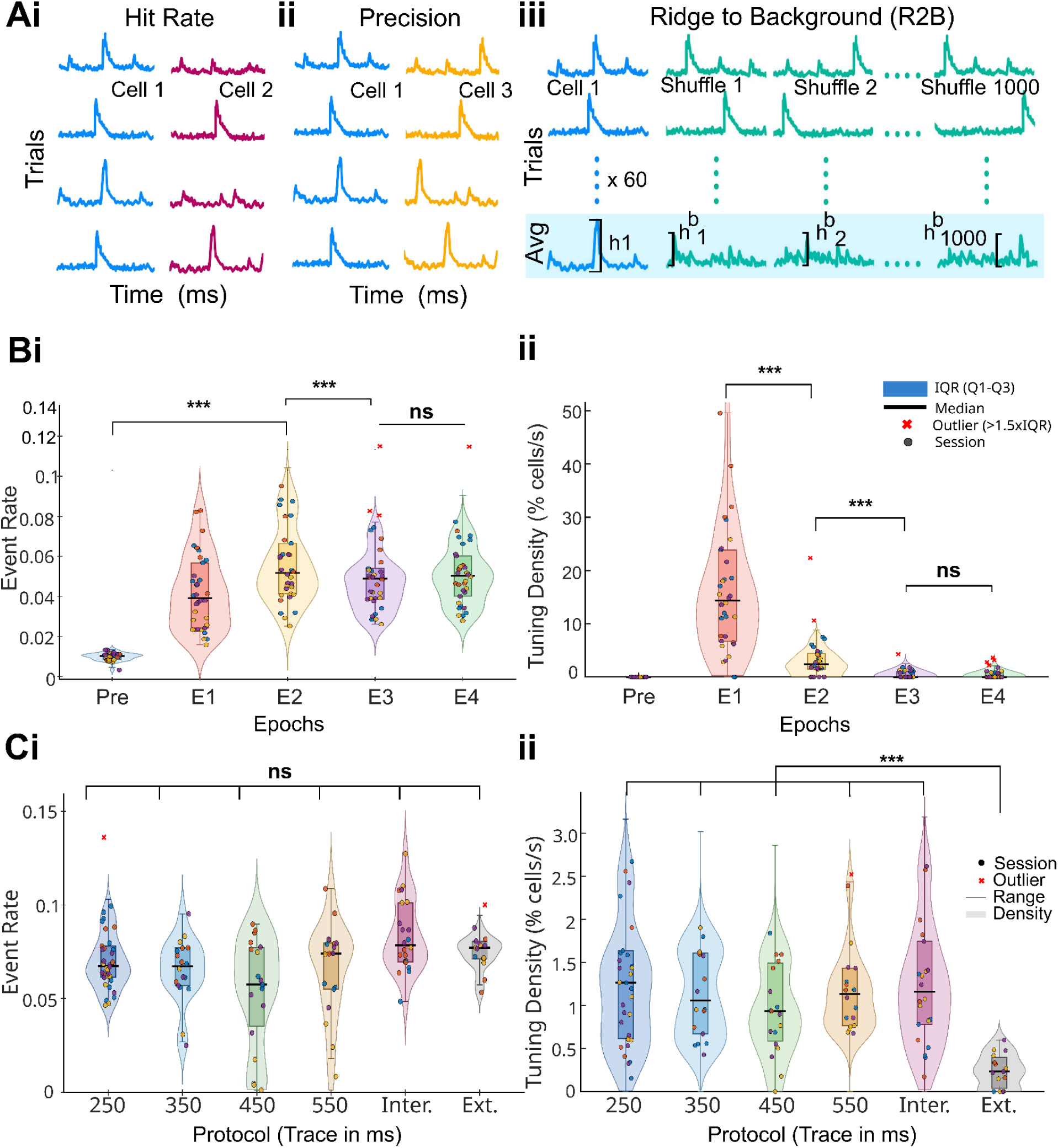
Time cell activity over the course of learning. **A:** Schematic of time cell identification criteria. Left: Hit rate — fraction of trials with an event within the target window. Middle: Precision — consistency of event peak timing across trials. Right: Ridge-to-background (R2B) ratio — observed peak compared to shuffled bootstrap data. **B:** Population event rate (Bi) and time cell tuning density (Bii) across temporal epochs (SoAn1, 250ms protocol; n=34 sessions, 4 mice). **Bi:** Event rate per epoch (fraction of cell-frames exceeding 2 SD above per-cell pre-CS baseline). Brackets: selected bootstrap comparisons (10,000 iterations); all pairwise epoch comparisons significant (p≤0.009) except E3 vs E4 (ns, p=0.197). Friedman (n=4 mice): χ²(4)=13.40, p=0.0095. **Bii:** Tuning density (%/s) per epoch. Bootstrap: E1>E2 (p<0.001), E2>E3 (p<0.001); E3 vs E4 (ns, p=0.77). Friedman (n=4 mice): χ²(4)=14.8, p=0.005. **C:** Cross-protocol comparisons (n=126 sessions total, 4 mice). **Ci:** Population event rate pooled over the post-CS window; no significant protocol difference (Friedman, n=4 mice; χ²(5)=6.86, p=0.232). **Cii:** Time cell tuning density (pooled post-CS window, %/s) across protocols (n=126 sessions, 4 mice). Omnibus Friedman test (n=4 mice): χ²(5)=9.57, p=0.088, ns. Bracket: bootstrap test comparing all conditioning sessions pooled (n=111) against Extinction (n=15); observed difference = 1.08 %/s (95% CI [0.88, 1.30]), bootstrap p<0.001 (N=10,000 iterations), Cohen’s d=1.13. Extinction showed consistently lower density than all conditioning protocols (0.24±0.20 %/s vs 1.33±1.02 %/s); **For all panels:** Violin shapes: KDE of session-level values (non-outlier sessions). Horizontal lines: median; boxes: IQR; whiskers: 1.5×IQR. Dots: individual sessions coloured by animal. Red crosses: upper outliers (>Q3+1.5×IQR), shown at true values and retained in all analyses.

As previously reported, sensory stimuli trigger activity in the hippocampus (Figure 3Bi). Population event rate was significantly modulated across epochs during paired conditioning (Friedman, n=4 mice; χ²(4)=13.40, p=0.0095; Figure 3Bi). Activity was low and tightly distributed during the pre-CS baseline (Pre: 0.030±0.002), rose sharply following CS onset through E1 (0.062±0.019) to peak at E2 (0.077±0.023), then declined to a stable plateau at E3 (0.070±0.019) and E4 (0.072±0.017; all values fraction of suprathreshold cell-frames). All pairwise epoch comparisons were significant (bootstrap p≤0.009) except E3 vs E4 (p=0.197), confirming a post-CS peak in population-wide recruitment that stabilises after the trace interval. The Bonferroni-confirmed contrast was Pre vs E2 (p=0.018).

To examine how time-cells distribute over the course of a trial, we computed tuning density, defined as the fraction of cells showing time-tuned activity per 1-second window. Within the 250ms protocol, tuning density was highest immediately following CS onset and declined steeply across successive epochs (Friedman, n=4 mice; χ²(4)=14.8, p=0.005; Figure 3Bii). E1 showed the highest density (8.66±7.39 %/s), significantly greater than E2 (1.70±2.10 %/s; bootstrap p<0.001), which in turn was significantly greater than E3 (0.26±0.45 %/s; bootstrap p<0.001). E3 and E4 (0.29±0.50 %/s) did not differ significantly (bootstrap p=0.77), indicating that temporal representation drops sharply within the first three seconds after CS onset and plateaus thereafter.

Neither population event rate nor time cell tuning density varied significantly across training protocols. Event rates were consistent across all protocols (Friedman, n=4 mice; χ²(5)=6.86, p=0.232; Figure 3Ci). Tuning density did not differ significantly across conditioning protocols (Time cell peaks pooled across the full post-CS window, Friedman, n=4 mice; χ²(5)=9.57, p=0.088; Figure 3Cii). However, Extinction showed markedly lower density than all conditioning protocols (0.24±0.20 %/s vs 1.33±1.02 %/s, p =0.0002; Extinction sessions (n=15) compared against all conditioning sessions (n=111, pooled across all 5 protocols), bootstrap p<0.001, N=10,000 iterations; Cohen’s d=1.13; Mann-Whitney U p<0.001). In addition to tuning density, the ratio of time cells to active cells also did not change with learning (Supplementary Figure 2). Epoch-resolved analyses reveal that protocol differences are localised specifically to early post-CS epochs rather than distributed uniformly across the full window (Supplementary Figure 3). To obtain a finer comparison between pre-learning with later stages, we compared session 1 for all animals with the day of criterion and last day of pairing for the 250ms sessions, and again did not find any dependence (Supplementary Figure 4)

### Time cells retain their tuning in the extended sequence despite changes in trace interval

Next we performed the crucial test of comparing time-cell tuning between the 250ms and 550ms blocks of the interleaved trials. Previous studies using working memory tasks with different delays found some consistently tuned cells, but many cells changed their tuning or were newly recruited (MacDonald 2011).

As a control, we first tested if there was an underlying tuning in timing of peak activity of all cells including non-time cells. (Figure 4A). We sorted cells by time of activity peak for 250ms trials, and tested whether the activity time-series for the 550ms trials were correlated (Methods). No significant correlation was observed (Table 1, Figure 4A). We also compared correlations for the probe trials, and again did not observe (Figure 4Aiii, iv). We repeated this by sorting first according to peaks for 550ms trials and testing for the 250ms trials (Figure 4B). As before, there was no peak ordering (Table 1). Thus, as expected for non-time-tuned activity, simple peak-time sorting does not yield a correspondence between the two blocks.

**Fig. 4.**
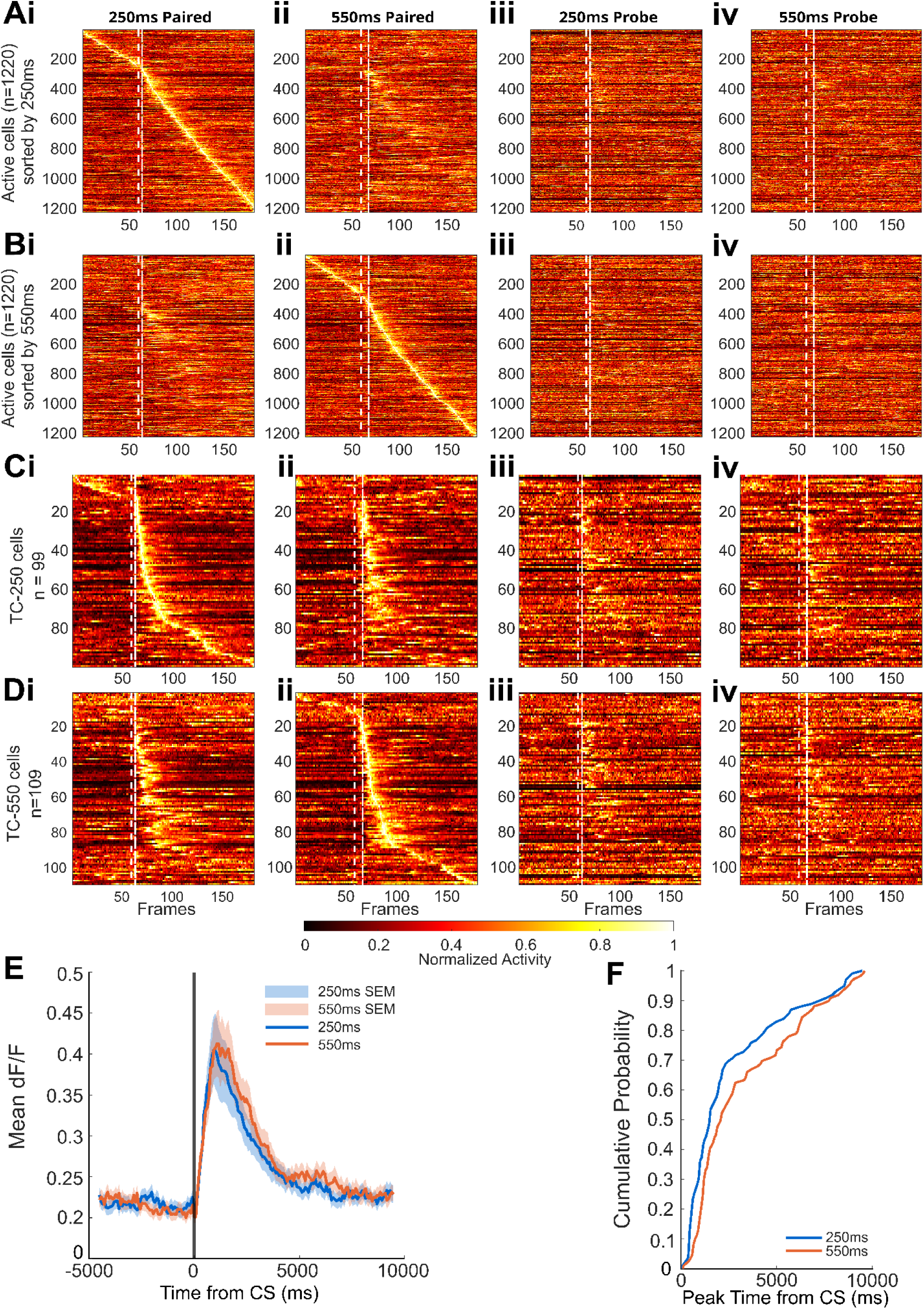
Time cell tuning is independent of trace interval. **A-D:** Activity heatmaps for all cells for interleaved trials showing normalised fluorescence (colour scale: 0–1) for each cell (y-axis) across frames (x-axis, 181 frames total). CS is indicated by spaced white dashes and US indicated by the dense white dashes. A and C are sorted per the peak timing of the 250ms trials and B and D as per 550 ms trials (B,D). The four columns in each row correspond to: (i) paired 250 ms trials, (ii) paired 550 ms trials, (iii) 250 ms probe trials, and (iv) 550 ms probe trials. Cells were sorted by peak activity time as described below. **A:** All active cells (n = 1220), sorted by peak time in paired 250 ms trials (Ai). A clear sequential diagonal is visible only in Ai, confirming the sort order. No corresponding diagonal appears in Aii, Aiii, or Aiv, indicating that peak-time ordering for non-time-tuned active cells does not generalise across conditions (statistics in Table 1). **B:** same as A, sorted as by peak time in paired 550 ms trials. The diagonal is visible only in Bii, with no ordering preserved in Bi, Biii, or Biv, **C:** Similar to A, but only considering time cells identified in the 250ms condition (n=99). Sequential peak-time ordering is preserved across all four conditions (Ci–Civ), indicating that time cell tuning is maintained when the trace interval changes and during probe trials. The diagonal is visible up to approximately frame 100 (∼3 s after CS). Probe trial heatmaps (Ciii, Civ) show a fainter diagonal because probe trials constitute only ∼10% of total trials, reducing the signal-to-noise in the trial-averaged trace (statistics in Table 1). **D:** Similar to C, but for 550 ms trials. As in C, sequential peak-time ordering is preserved across all four conditions, confirming bidirectional stability of time cell tuning. **E:** Mean dF/F activity trace (± SEM, shaded) for TC-250 (blue) and TC-550 (orange). **F:** Cumulative probability plot for peak positions. Peak time distributions did not differ significantly between 250ms and 550ms trace conditions (Mann-Whitney p = 0.0542; Kolmogorov-Smirnov: D = 0.142, p = 0.2247).

**Table 1:**
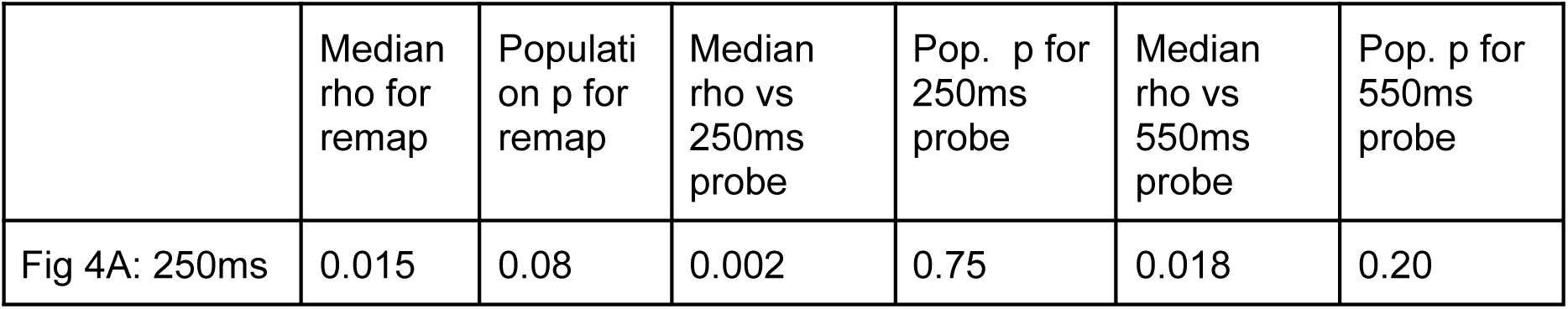
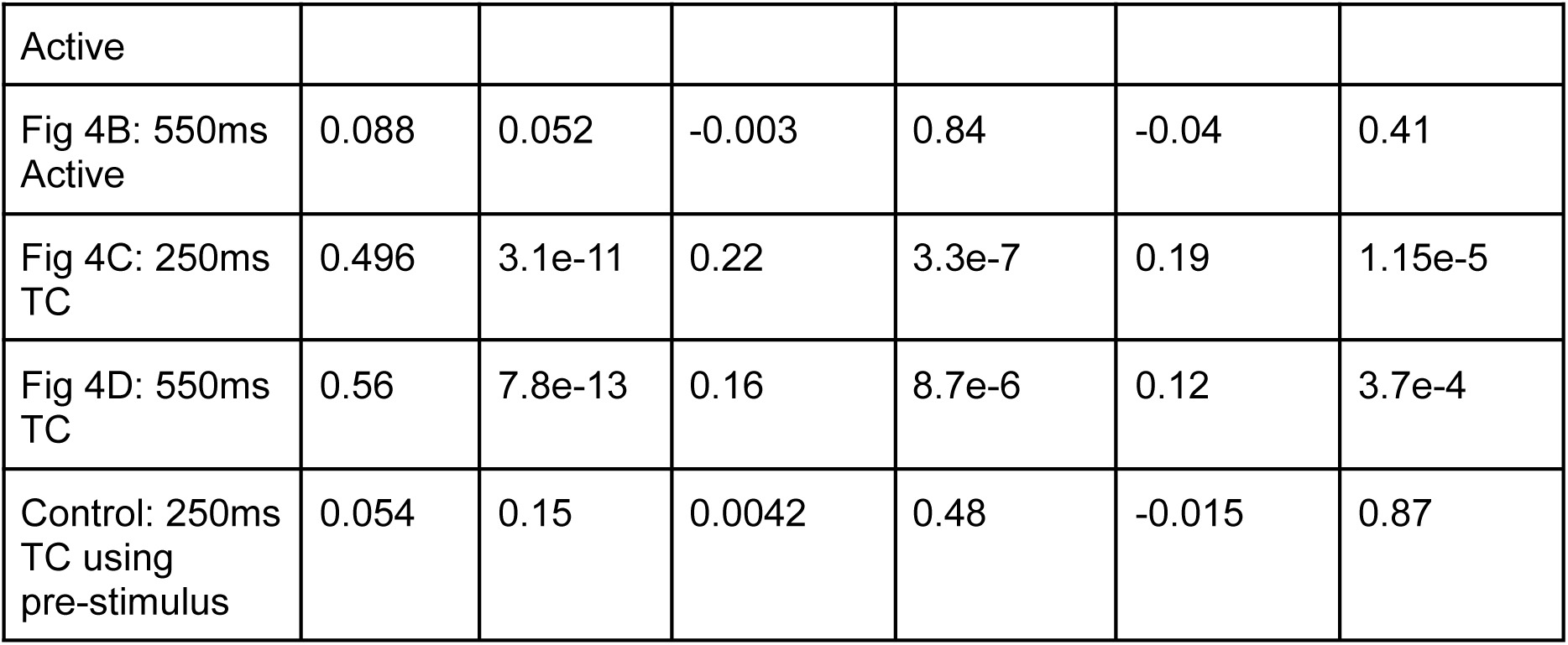
Time-course of post-stimulus activity is not correlated for Active, non-Time cells, but strongly correlated across different trace intervals and against probe trials for Time Cells.

We then tested if time cells differ in their tuning when representing different trace intervals (Figure 4C, D). As for the Active cell controls, we compared the activity time-course from CS to CS+3s for time cells identified in the 250ms condition (TC-250, Figure 4Ci) with the same cells during the three other conditions: 550ms sessions (’Remap’, Figure 4Cii), 250ms probe (Figure 4Ciii), and 550ms probe (Figure 4Civ). We used Spearman’s test and aggregated the rho values for the population of cells using the Wilcoxon signed-rank test to obtain a population significance level for correlation (methods). All the cases were strongly correlated (Table 1). We performed a similar analysis using time-cells sorted as per 550ms trials (Figure 4D), and again, all cases were strongly correlated (Table 1). As a further control we performed the same calculations as in 4C, but using the 3 seconds preceding CS, and found that none of the cases were significantly correlated (Table 1).

The behavioral data showed that the maximum eyeblink was later in the 550 ms trials (Fig 2). We asked if the peak time for time cells was also delayed. First, we simply plotted the distribution of time-cell activity as a function of time following CS. The peaks of the 550 ms trials were slightly broader (Figure 4E). To test if this was a significant shift, we plotted the cumulative distribution of the 250 and 550ms trial activity peaks (Figure 4F). They did not significantly differ between conditions (Mann-Whitney p = 0.0542; Kolmogorov-Smirnov: D = 0.142, p = 0.2247).

Together, this analysis on the interleaved trials showed two things: time cells do not remap when the trace interval is changed between 250ms and 550ms, and further, the time course of activity is similar even when the US is missing altogether in the probe trials. These results support an *absolute timing* coding scheme, where cells encode specific durations (e.g., ’2 seconds’) rather than rescaling to the total interval length.

### Time cell timing profiles vary with epoch and trace interval

A striking feature of our time-cell activity was that time-tuned activity extended much beyond the trace period. Specifically, even for our 250 ms trace period there was time cell activity out to ∼9 seconds after the CS (Fig. 4C,D). Post-stimulus time-cell activity has also been reported for ∼1.5 seconds by Bhattacharjee et al.

As a first-order examination of the epoch structure, we asked if time-cells were more common in early epochs. We found that the first two epochs contained similar proportions of time cells (Epoch 1 (0-1s): 34.6%, Epoch 2 (1-3s): 35.1%), and then the number declined steeply (Epoch 3 (3-6s): 15.9%, >6s: 14.4%) (Figure 5A). The 250 ms condition had a higher proportion of cells in Epoch 1 (42.4% vs 27.5%, χ²(1), p = 0.024; session-level bootstrap p = 0.009), while the 550 ms condition showed enrichment in Epoch 2 (43.1% vs 26.3%, χ²(1), p = 0.011; session-level bootstrap p < 0.001), consistent with a shift in temporal coverage toward the respective reinforcement times. No significant differences were observed in later epochs (p > 0.10).

**Fig. 5:**
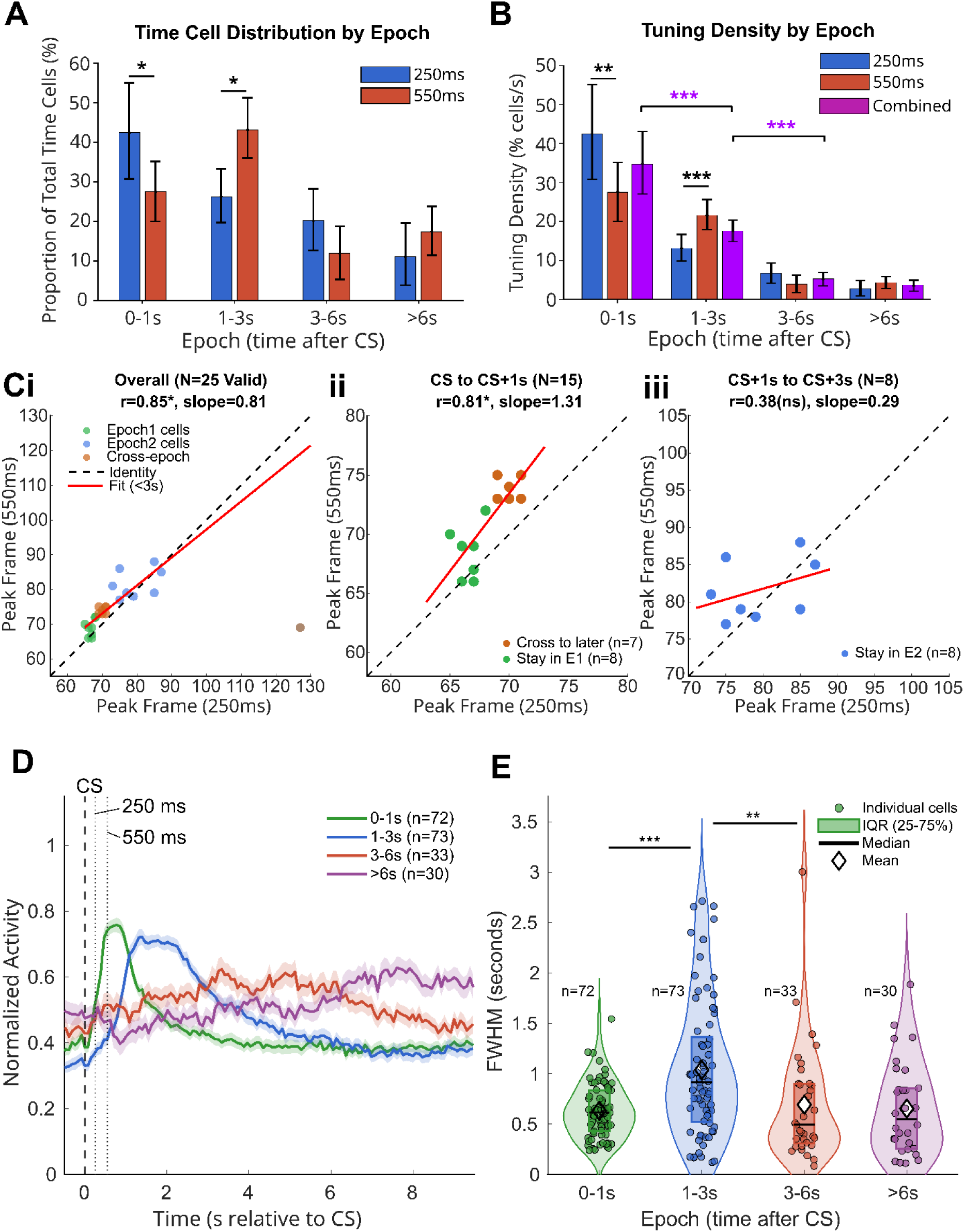
Time cell distributions and peak widths vary with TEC interval and trial epoch. **A:** Time cell distribution by epoch for 250 ms and 550 ms conditions separately. Error bars represent 95% confidence intervals from session-level bootstrapping (10,000 iterations). Significance brackets indicate between-condition differences (session-level bootstrap, two-sided). *p < 0.05. **B:** Tuning density of time cells (%/s) decreases over trial epochs for 250 ms, 550 ms, and combined conditions (Cochran-Armitage trend test, T = −11.89, p < 0.001 for combined). Brackets indicate significant adjacent-epoch decreases in the combined condition. Error bars represent 95% confidence intervals from session-level bootstrapping (10,000 iterations): sessions were resampled with replacement and their constituent cells pooled to recompute density, accounting for the hierarchical structure of cells nested within sessions. Significance: *p < 0.05, **p < 0.01, ***p < 0.001. **C:** Peak times are aligned between 250ms and 550ms trials, using linear regression comparisons. **Ci:** All epochs (N = 25). **Cii:** Epoch 1, CS to CS+1s (N = 15). **Ciii:** Epoch 2, CS+1s to CS+3s (N = 8). **D:** Mean normalised activity traces for cells grouped by peak epoch (combined conditions; n values as indicated). **E:** Violin plot of tuning width (FWHM) distributions by epoch. Cells peaking in the second (1-3s) epoch have significantly broader tuning than those in epoch 1 (Mann-Whitney U, p < 0.001), and in epoch 3 (p = 0.005).

To better characterize time-cell abundance, we computed tuning density, defined as the percentage of time cells firing in each second of an epoch, with units of %/s (Figure 5B). Time cell tuning density decreased monotonically across temporal epochs (Cochran-Armitage trend test, T = −11.89, p < 0.001; Fig. 5B). Combined across both trace conditions (N = 208 time cells, 22 sessions, 4 mice), density was highest in the first second after CS onset (34.6%/s, 95% CI: 26.6–42.5) and declined approximately 10-fold by 6 seconds (>6s: 3.6%/s, CI: 2.3–5.1). Adjacent-epoch decreases were significant from epoch 1 to epoch 2 (p < 0.001) and from 1-3s to 3-6s (p < 0.001), with density reaching a plateau beyond 3 seconds (3-6s vs >6s: p = 0.093). The distribution of time cells across epochs was significantly non-uniform for both trace conditions (χ²(3) > 88, p < 10⁻¹^8^). In other words, time was most densely encoded in the immediate aftermath of the stimulus, but reliable time-cell activity persisted well beyond apparent stimulus salience.

Do time cells have the same time preference in 250 vs 550 ms trials? In Figure 4 we had found that the time-courses of post-CS activity were strongly correlated. Here, we tested if peak times were linearly related. We plotted the time of peak activity for cells which passed the time-cell criterion for both 550 and 250 ms trials, for different combinations of epochs (Figure 5C). The slope was nearly 1 for epoch 1+epoch 2 combined (r = 0.848, slope = 0.81 ± 0.11, p = 0.093 ns), and for epoch 1 alone (r = 0.811, slope = 1.31 ± 0.26, p = 0.254). Epoch 2 showed a slight but significant deviation from a slope of 1, though this may reflect the small sample (n = 8, r = 0.379 ns, slope = 0.29 ± 0.29, p = 0.049). Thus time tuning is nearly maintained between the two trial durations.

Given that time cell tuning propensity varied with epoch, we next asked if the tuning width also varied. We computed full width at half maximum (FWHM) for time cells in each of these epochs and found that tuning width varied significantly across epochs (Kruskal-Wallis H = 17.98, p = 0.0004). Post-hoc pairwise comparisons (Mann-Whitney U) revealed that cells peaking in the second (1-3s) epoch had significantly broader tuning than all other epochs (vs epoch 1: p < 0.001; vs epoch 3: p = 0.005; vs epoch 4: p = 0.007). Epochs 1, 3 and 4 did not differ from one another (all p > 0.44).

In summary, we identified at least three distinct epochs for time-cell responses to our trace-conditioning protocol. Time-cell tuning density declined with successive epochs, but tuning width was greatest in the second epoch.

### Time cells decrease following behavioural extinction

The final stage of our behaviour protocol was to introduce behavioural extinction by running sessions with CS but without US reinforcement. This leads to a decrease in performance (eye-blink prior to US time) which seems to mirror the initial rise in performance during the first learning phase before reaching the 60% criteria (see Fig.2: A). To test whether hippocampal time cells show similar dynamics during the prelearning and extinction phase, we compared neural representations between early learning (pre-criterion 250ms sessions, n=18) and extinction (sessions, n=15). Notably, even as the overall activity, calculated as calcium event rates, were slightly elevated during extinction (Figure 6A) (0.062±0.008 vs 0.055±0.011, p=0.032), the time cell proportions were reduced during extinction compared to pre-criterion sessions (Figure 6 B) (R2B: 13.7±10.6% vs 2.4±2.0%, p<0.0001, Cliff’s d=0.88; TI: 7.3±6.2% vs 1.1±1.9%, p=0.0002, Cliff’s d=0.76). There was no apparent dependence of time cell fraction upon behavioural performance (Figure 6C), nor were the number of recorded cells significantly different (Figure 6D). These results suggest that the neural representation of extinction is fundamentally different from reversion to a pre-learning state (Bouton, 2004; Myers & Davis, 2007). We interpret this to mean that there may be an active process of removal of time cells and, consequently, of this potential neural activity linkage from CS to US.

**Fig 6:**
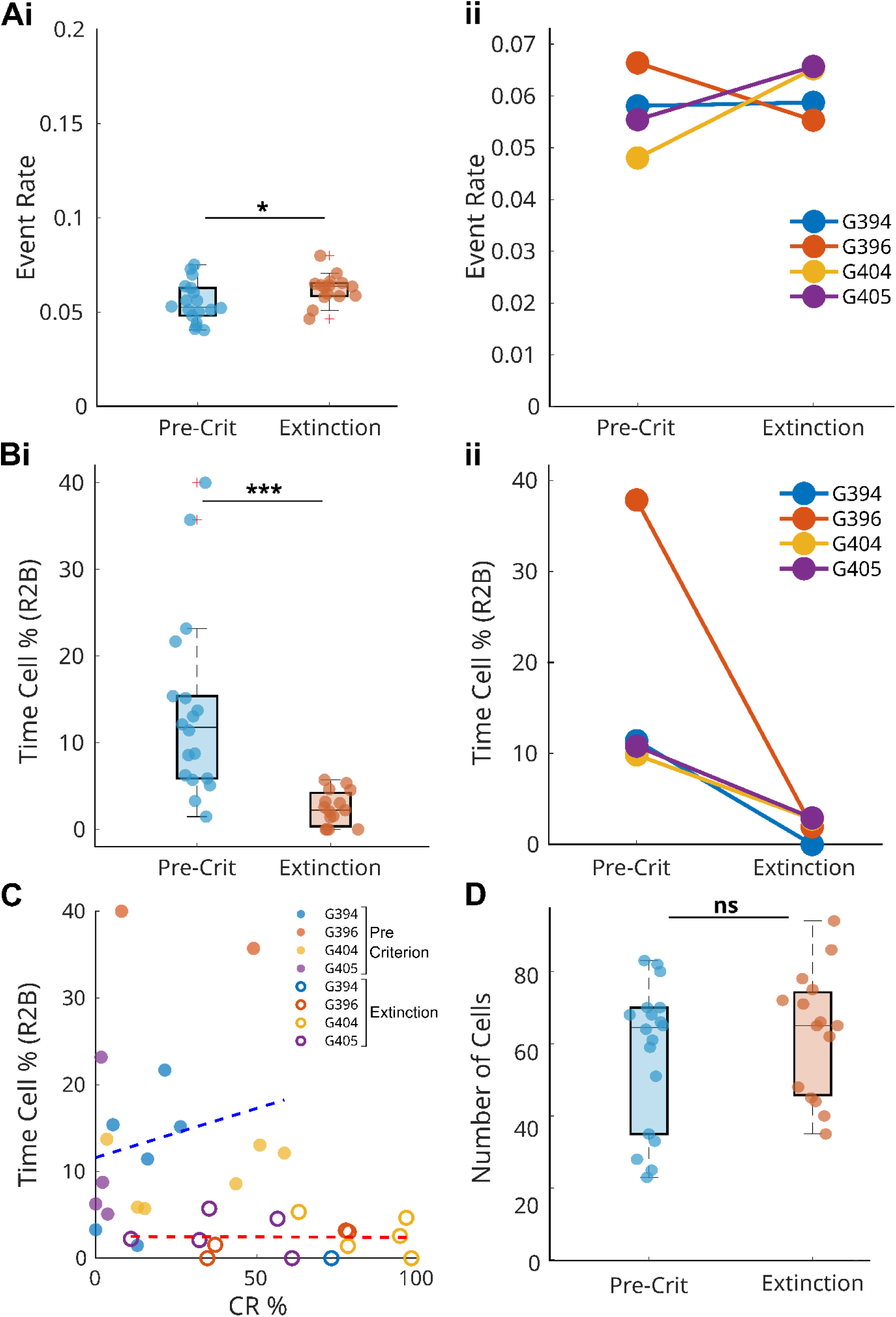
The proportion of time cells decreases during behavioural extinction. **A:** Controls to show that the event rate over all cells increases slightly when pre-learning is compared to extinction (Mann-Whitney U: p=0.032*, Cliff’s d = -0.444, effect size medium) with (Aii) 3 of the 4 animals showing small but general increase in event rate during extinction. **B:** (i)Time cell percentage drops steeply between pre-learning and extinction phases. (Mann-Whitney U: p< 0.0001*, Cliff’s d = 0.878, very large) and (Bii) per-animal trend also shows a stark decrease in TCs.**C:** Time cell proportions do not correlate with behavioural performance expressed as correct response percent, either for pre-learning (Filled circles, Spearmans r=0.25, p= 0.325) or during extinction (Open circles, Spearman’s r=0.02, p=0.949). **D:** There is no significant difference in the number of recorded cells between the two cases, thus ruling out technical confounds related to number of cells compared (Mann-Whitney U: p = 0.4156; Effect size (Cliff’s d): -0.170).

**Fig 7:**
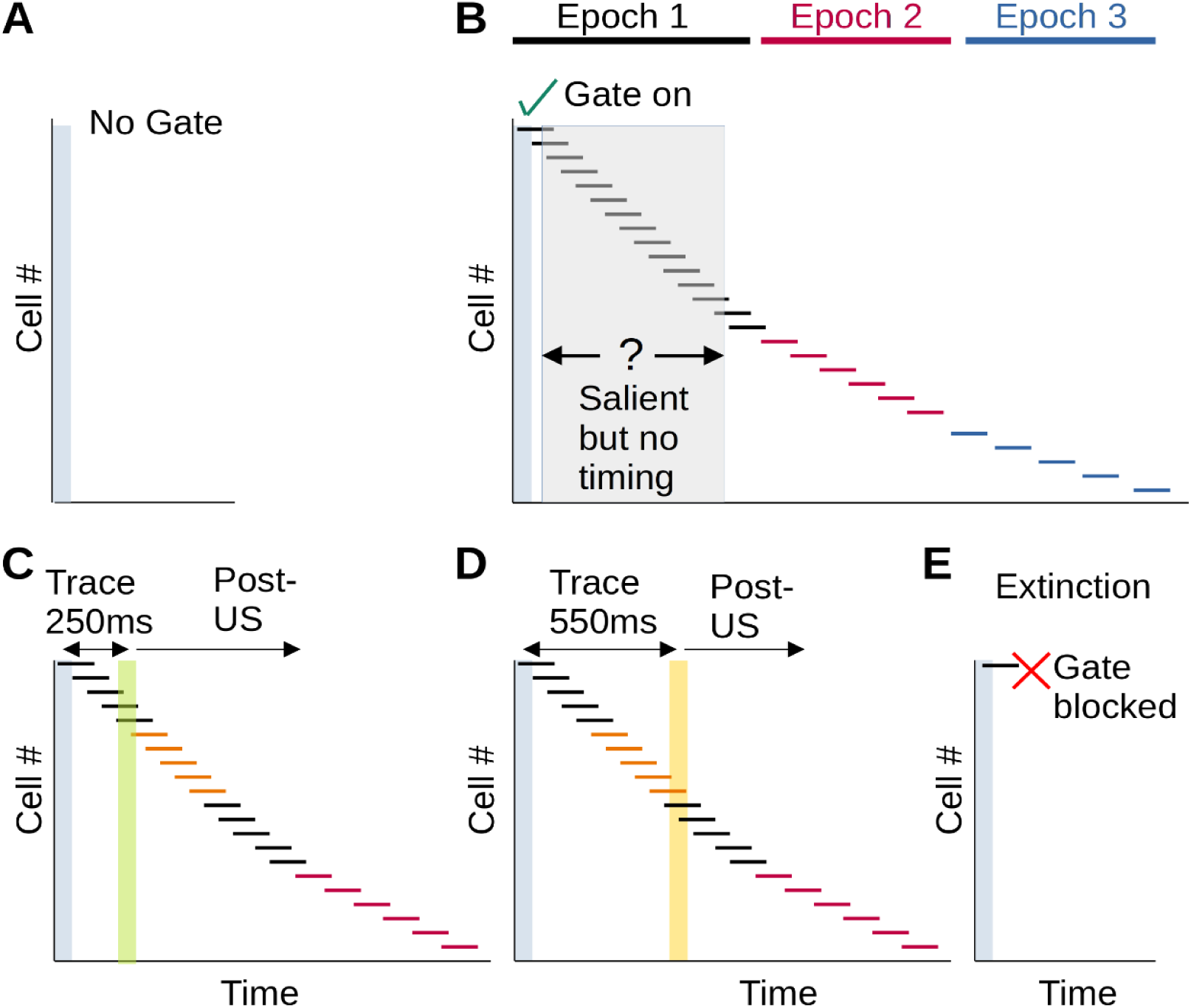
Hypothesis and life-cycle of time-cells. In all panels the grey vertical bar represents the conditioned stimulus. **A:** CS without any salience. No gating has yet occurred to bind this to a time-cell sequence. **B:** CS with salience gates the time cell sequence, even though US timing has not yet been learnt. Note multiple epochs of time-cell activity. **C:** CS followed by US (green bar) after 250 ms. The same time-cell sequence as in B is active, only the first two epochs shown. **D:** CS followed by US (yellow bar) with a trace interval of 550 ms. All time cells retain their time tuning. Specifically, the cells indicated in orange had been after the US in panel C, but are now before the US. Nevertheless they do not change their tuning. **E:** Extinction differs from panel A in that there is a latent sequence which is now actively blocked.

## Discussion

We examined time cell tuning in a TEC paradigm with a range of trace intervals, including a set of sessions where the trace interval was interleaved between 250ms and 550ms, and finally behavioural extinction. Time cells occur in the same fraction from the start of the learning process and during learning. Time cells are initiated with the CS, but there are at least three epochs following the CS in which their tuning density and peak widths vary, and time cells are observed up to 9 seconds post-stimulus. Remarkably, there is no remapping of time cells between the 250ms and 550ms trials, instead they remain on the same sequence until ∼3 seconds after CS. Behavioural extinction is accompanied by a steep reduction in the number of time cells.

### Time cell formation is independent of learning

A key departure point of our findings from previous work in working memory tasks (MacDonald et al., 2011, 2013) is that we observe time cell formation independent of learning. In previous work using working memory tasks, it has been shown that the time cells appear in sync with behavioural outcomes (Gill et al., 2011; Taxidis et al., 2020). In these previous studies, the time-scales involved (∼10 seconds) are more than an order of magnitude longer than the trace intervals of 250ms to 550ms which we employ in TEC. Furthermore, the task demands differ, since in TEC the animal has to maintain precise timing information to pre-empt the arrival of the air-puff. Thus it seems plausible that the underlying biophysical and network mechanisms may differ between TEC and working memory, though the computational outcome of time-cells tiling an interval is remarkably consistent.

The relevance of time cells in working memory has been challenged by studies which report that working memory for tasks longer than ∼5 seconds does not depend on the presence of time cells (Yuan et al., 2025). While the differences with our TEC protocols do not allow us to directly comment on this debate, it is intriguing that we find strong post-stimulus time cell activity up to at least 3 seconds (Figure 4). We speculate that this post-stimulus time cell trace may be a bridge between the sub-second time-scales of TEC and the multi-second time-course of working memory tasks.

A more direct comparison is with earlier work on a TEC task by Modi et al., (2014) where time-cell formation was linked to the emergence of conditioning within a single session. In recent work using some of the same data, we suggest that the intensity of the stimulus in Modi et al. may account for the accelerated learning (Bhattacharjee et al., 2024).

### Time cells do not remap with trace interval

A core finding from our study is that time cell tuning in TEC is consistent regardless of trace interval. This contrasts with previous findings for working memory tasks, where a few time cells retained their tuning, but linear stretching and remapping were more common (MacDonald et al., 2011). We suggest there are methodological as well as conceptual differences which may account for this divergence of findings. On the methodological front, we designed our experiments to eliminate day-to-day learning and alternated the long and short trace period blocks within <300 seconds of each other. This is a more compact protocol than used in previous studies (MacDonald et al., 2013), in which there were three distinct time blocks during the session. The training paradigms also differed, ours being TEC and Macdonald et al using a working memory task involving object-odor association. Conceptually, we suggest that the time-cell formation mechanisms at the 250ms time-scale may be fundamentally different from those for ∼10 seconds. Let us call these ‘fast’ and ‘slow’ time-cell mechanisms. Fast mechanisms are easier to relate to timescales of cellular biophysics (Tsodyks & Markram, 1997), and to network mechanisms such as synfire-chains (Diesmann et al., 1999; Kumar et al., 2008, 2010). Slower timescale sequences may emerge from signaling mechanisms (Bhalla, 2002) or traversal through a series of attractors in recurrent networks (Buonomano & Laje, 2010; Buonomano & Maass, 2009; Laje & Buonomano, 2013; Rajan et al., 2016). This division of time-scales suggests a possible analogy with place-cell sequences, which are slow during locomotion (seconds time scales) and fast during replay (100ms time-scales)(O’Keefe & Recce, 1993; Skaggs et al., 1996). A clue to the possible upper limit of the ‘fast’ mechanism comes from our observed period of complete overlap of up to CS+3 seconds. Interestingly, Yuan et al. also report that reliable time cells are only present in the first few seconds of a working memory task (Yuan et al., 2025).

Another point of reference for remapping of hippocampal neuronal representations is when the context changes (Bostock et al., 1991; Leutgeb et al., 2005; Muller & Kubie, 1987). In a similar TEC protocol to our current work, Bhattacharjee et al. (Bhattacharjee et al., 2024) changed the context between visual and auditory stimuli, presented in interleaved blocks. This previous study showed that most time cells were indeed modality dependent. Similar modality-dependent time-cells have been reported for working memory time-cells (Taxidis et al., 2020). Thus remapping can occur for TEC time cells due to modality, but does not occur due to changes in task interval. Together, these observations suggest that trace interval changes are handled by the network in a distinct way from context changes, and do not undergo remapping even though the network does provide a substrate for it.

### The extended time-cell trace may act as a timing template

Previous studies have proposed that intrinsic brain activity sequences may act as templates for correlating events with a consistent timing structure (Azizi et al., 2013; Vaz et al., 2023). For example, Dragoi et al. have suggested that ‘preplay’ sequences may play such a role in subsequent recruitment of place cells (Dragoi & Tonegawa, 2011; Eichenbaum, 2014). Our study provides two lines of evidence to support this. First, as discussed above, we find that the time-cell sequence starts with the conditioned stimulus, and even though the US is a much stronger stimulus in terms of salience and triggering network activity in naive animals (Berger et al., 1976, 1983; McEchron & Disterhoft, 1997), it continues with the same sequence regardless of the trace interval (Figure 4). Second, we analyzed time cell sequences in probe trials which lack the US, and found that these were highly correlated with the sequences following regular CS-US pairing (Figure 4). Thus in both cases the weaker light stimulus triggered a sequence which was not altered despite timing shifts, or even absence of the much stronger air-puff stimulus.

### Behavioural extinction leads to time-cell elimination

The extinction of a behaviour is typically understood as a process of learning not to respond, rather than simply forgetting. For example, in fear conditioning, Chattarji et al have shown that fear learning can be extinguished by exposure provided the fear learning happens before but not after a stressful experience (Rahman et al., 2018). Additionally, when extinction is obtained in fear or operant conditioning, it usually only takes one or a few reinforced trials for the original behaviour to be completely retrieved (Bouton, 2004; Bouton & Bolles, 1979; Rescorla, 2004). We suggest that our observations of the removal of time cells during extinction also fall within the framework of an active process to turn off the time cells. Two of our observations support this interpretation. First, we see time cells even before the animal has formed a behavioural association (Sup Fig:2; Bhattacharjee et al., 2024). Thus, if all that was happening in extinction was to forget the association, it would be a reversion to the pre-learning condition and we would expect the time cells to remain. Second, time cells do not change tuning during probe trials, where reinforcement is also missing. Specifically, the tuning is intact even after the US would normally have occurred. Thus the loss of time cells following extinction is both different from baseline responses before learning, and from pre-extinction responses to CS alone. There is supporting evidence from lesions (Clark et al., 2002; Moyer et al., 1990; Solomon et al., 1986) and activity blockage (Bigus et al., 2024; Dias et al., 2021) that time-cell activity is likely to be causal in learning of timed events, but see (Ahmed et al., 2020; Sabariego et al., 2019). We speculate that time cells are indeed causal in eliciting the trace eyeblink response, and the network actively suppresses these time cells as part of learning to ignore the conditioned stimulus as a signal for a forthcoming eye-puff.

### Hypothesis: an extended time-cell sequence templates activity over multiple trace intervals

Based on our observations, we propose a simple model for time-cell function tuning for TEC tasks, which markedly differs from what is seen in operant, working-memory, and treadmill-running tasks (Gill et al., 2011; Kraus et al., 2013; Lee et al., 2006; MacDonald et al., 2011). We propose that time cells form as soon as an animal regards a stimulus as salient, but even before the precise timing relationship of stimulus to response is learnt (Figure 1-5). This interpretation has close parallels with place-cell formation on first exposure to a novel spatial environment (Frank et al., 2004; Hill, 1978; Wilson & McNaughton, 1993). This time-cell sequence extends for a few seconds, without being bounded by the US, since at this stage the timing of the US relationship has not even been learnt. When behavioural conditioning occurs, the US events map onto this existing sequence, and when there are changes in the timing relationship, this same time-cell sequence is re-used, without remapping or stretching of time-cells. There are two corollaries of this model. First, we posit that there is an epoch structure of time cells in trace conditioning, and it is only in the first (CS to CS+1 second) epoch where there are high densities of time cells, and eyeblink conditioning is readily acquired. This may account for the observed behavioral limitation on the trace interval which mice can readily learn (<750 ms, (Kishimoto et al., 2001; Tseng et al., 2004)). Such an epoch structure differs from the logarithmic representation of time proposed by Cao et al. (2022), possibly because of the 10x longer time-domains of the earlier analysis. This leads to the prediction that the first, high TC density epoch should be of greater duration in other model organisms such as rats, which can form longer associations (McEchron et al., 1999; McEchron & Disterhoft, 1997; Weiss et al., 1999). Second, we propose that there is a crucial upstream gating step to link the initial CS to a latent time-cell sequence. Such latent sequences have been proposed in place-cell sequences (Dragoi & Tonegawa, 2011)and as general templates for linking discontiguous events(Villette et al., 2015). This gating is activated early in learning when CS salience is learnt, even before the time relationship to a US. We envision this as similar to the nearly single-trial learning of place-cell representations. The gating can also be disabled, as an active process leading to extinction.

This model differs from previous studies in several key respects. First, it discusses events on time-scales about an order of magnitude faster than object-odor pairing time-cell experiments (MacDonald et al., 2011, 2013). Second, it proposes invariance of time-cells within the first two epochs regardless of trace interval, unlike adaptive time-cells in variable-duration working memory tasks (MacDonald et al., 2011). Third, it proposes a role for post-stimulus time-cell activity as a sequence template to which novel or delayed stimuli can be associated. Fourth, it explicitly links salient events to latent time-cell sequences through a gating process. Finally, it posits multiple epochs in a time-cell sequence, of which the fastest underlies trace-conditioning, and the slowest may be a bridge to the ∼10 second time-scale of many studies with working memory or treadmill running intervals.

## Methods

### Animals

In vivo two-photon calcium imaging studies utilized 4 adult mice (12-24 weeks old) expressing GCaMP6f (GCaMP6f+/-; C57BL/6J-Tg(Thy1-GCaMP6f)GP5.17Dkim/J × C57BL/6J). Behavioral experiments without calcium imaging employed an additional 4 GCaMP6f-/- mice of the same age range. The mice were sourced from The Jackson Laboratory, which were then bred and housed in the NCBS animal house. Animals group-housed (2-4 per cage) under a 12-hour light/dark cycle.

### Surgical Procedures

A cranial window was implanted to access the hippocampus in a protocol similar to that used by Dombeck et al., 2010. Among the four mice implanted for imaging, two were implanted on the right hemisphere, and the other two on the left hemisphere. Animals were on water restriction for 3-5 days pre-surgery, while ensuring the animals retained 80% baseline body weight. Isoflurane anesthesia (ISIFRANE 250, Abbott, North Chicago, IL, USA) was delivered via carbogen (95% O₂, 5% CO₂) using a tabletop vaporizer system (VetEquip Inc., Items 901801 and 911103). Vapor flow rates of 2.5-3 L/min during induction and 1.2-1.5 L/min during maintenance were employed. Core temperature was regulated using a temperature stabilized heating pad (TC-1000 Temperature Controller with heating pad, CWE Inc., USA). Head stabilization utilized cheek clamps (Mouse Stereotaxic Adaptor, Stoelting Co.), while chloramphenicol eye ointment (1% w/w) prevented corneal desiccation.

After trimming cranial fur and disinfecting the scalp with 70% ethanol, a circular incision exposed the skull from bregma to lambda, extending approximately 5 mm bilaterally to the sagittal suture. Cotton swabs and a bone scraper were used to remove the fascia and dry the skull surface. Reference marks in millimeters were placed on the skull. A custom stainless steel headbar (10 mm central aperture) was secured using UV-curing dental cement (3M ESPE RelyX U200, Shade: TR) without skull screws.

A dental drill created a 3 mm diameter craniotomy centered at coordinates 1.5 mm left or right lateral and 2 mm rostral to bregma. Following dural removal with forceps, cortical tissue was aspirated using a blunted 26-gauge needle (26×1/2 DISPOVAN syringes, HMD Ltd.) connected to a vacuum line, pressure controlled using a roller clamp. Continuous irrigation with cortex buffer (125 mM NaCl, 5 mM KCl, 10 mM glucose, 10 mM HEPES, 2 mM CaCl₂, 2 mM MgCl₂, pH 7.35 adjusted with NaOH) prevented tissue desiccation. Bleeding was permitted for 5-10 seconds to facilitate clotting when necessary. Aspiration was done gradually over a period of 15-20 minutes to remove cortical tissue to a depth of 1 mm and to expose the corpus callosum fibers. Further aspiration was done to remove the superficial two layers of the corpus callosum while preserving the third layer close to the hippocampus.

Following buffer removal, the cavity was air-dried for 10-15 seconds until the glistening appearance disappeared. The hippocampus was overlaid with a minimal quantity of Kwik-Sil (low-toxicity silicone adhesive, World Precision Instruments, Inc.). A pre-assembled stainless steel cannula (3 mm outer diameter) was constructed with an affixed 3 mm coverslip (D263, CS-3R, #0 thickness, Warner Instruments, LLC) at its base using UV-curing glue (Norland Optical Adhesive NOA 81). The coverslip assembly was placed directly on the surface of the hippocampus, distributing Kwik-Sil evenly through its weight. This transparent adhesive layer minimized imaging-associated motion artifacts. Additional Kwik-Sil sealed gaps between the cannula exterior and craniotomy margins. Subsequent dental cement application covered the remaining exposed skull, flowed around the cannula’s lateral surface, and reinforced the headbar’s upper portion to minimize relative motion among headbar, skull, and cannula components.

Post-surgical recovery occurred following isoflurane cessation with continued carbogen delivery (1 LPM) until normal respiration resumed. Animals returned to home cages received 1 ml daily water throughout the experimental period, with ad libitum ibuprofen (2 ml/L) and enrofloxacin (1 ml/L) supplementation for 3-5 days.

### Experimental Setup

The apparatus employed a cylindrical foam treadmill (15.24 cm diameter, 11.43 cm length) with an axial metal rod enabling bi-directional locomotion (forward/backward). Headbar-implanted animals were secured via a custom clamp. All experiments occurred in darkness within a black anodized steel chamber housing the microscope. An in-house custom two-photon microscope was utilized for imaging.

Stimulus delivery systems included: a 480 nm blue LED positioned ∼7 mm from and centered to the snout for visual conditioned stimuli, and a wall-mounted carbogen supply (∼15 psi) regulated through a flowmeter (0.2-0.3 LPM, Cole Parmer) for unconditioned stimulus air delivery. The airpuff system consisted of clear PVC tubing terminating in an 18-gauge syringe positioned 5 mm from the left eye. Precise airflow timing was achieved using a solenoid valve (EV mouse valve, Clippard, Cincinnati, OH, USA). An additional roller clamp was used on the airflow line to provide finer control on pressure so as to optimize the pressure per animal until the blink response was reliable and reproducible and also ensuring the puff isn’t too harsh.

Eyeblink responses were captured at ∼200 fps using an infrared camera (Blackfly S USB3; BFS-U3-13Y3M-C, Teledyne FLIR LLC) connected via USB to a data acquisition laptop. An Arduino microcontroller coordinated LED, speaker, solenoid, and camera operations while providing TTL pulses for imaging synchronization. Custom Python software (https://github.com/BhallaLab/Mousebehaviour) controlled the behavioral apparatus.

### Behavioural Protocol during 2-P imaging

The trace eyeblink conditioning (TEC) paradigm, adapted from Siegel et al. (2015) incorporated a 50 ms conditioned stimulus (CS), variable trace interval, and 50 ms unconditioned stimulus (air-puff on the cornea). Six experimental stages with different trace intervals were implemented successively: 250ms, 350ms, 450ms, 550ms, Interleaved 250ms and 550 ms, and Extinction protocols. Each mouse went through all the behavioural protocols. Mouse 394 and 404 had a left hippocampal craniotomy, and Mouse 396 and 405 had a right-side craniotomy. Behavioural data for Mouse 404 was not considered because of light spillover from the scan laser to the eye, which obscured the eyeblink signal, however the animal went through the same sequence as other mice, of 7 days for 250ms training, followed by at least 4 days at each behavioural stage.

Handling and acclimation proceeded for ≥3 days until animals remained calm on the experimenter’s palm, coinciding with water restriction (∼1 ml per day per animal). This was followed by the surgical procedure. A 5-day post-surgical recovery period maintained 1 ml/day medicated water (antibiotic and analgesic), preserving body weight at 80% of pre-restriction baseline.

CS-US pairings were provided daily for approximately 60-trials per session (Figure1:Ai), with 10% trials pseudo-randomly designated as probe trials (CS-only). Upon achieving ∼ 60% conditioned response, the animals continued on the 250 ms protocol for an additional 2 days. This stabilized responses, after which they were shifted to the 350 ms trace interval protocol. On the day of switching the protocols, the mice went through two sessions with ∼ 30 trials each for both trace intervals independently. This was followed by ∼4-5 days of staying on the new trace interval (Figure1:Aii - iv). For 350 ms to 550 ms protocols, the switch was made based on observing consistent performance in the live behavior monitor.

Once the mice performed on independent trace interval pairings over multiple days, they were subjected to sessions with an interleaved block design, where on the same day and in the same session, the mice were exposed to both 250 ms and 550 ms trace pairings. The block structure consisted of consecutive 5 trials of 250 ms, followed by successive 5 trials of 550 ms, back to 5 trials of 250 ms, and so on. (Figure1:Av). The final stage of the behavior experiment was the extinction phase, where the mice were subjected to only the CS, and the air puff reinforcement was absent.

### Behaviour-Only Experiments

Four headbar-implanted mice underwent behavioral training without craniotomy, providing behavior only data without contributing to calcium imaging data. The behavior experiments were carried out following the same procedure and setting as that of the imaging animals.All four animals completed the six different stages of protocol learning similar to the imaging animals.

### In Vivo Two-Photon Imaging

Calcium imaging employed a custom galvo-scanning two-photon microscope (Model 6210H galvo, Cambridge Technology, Inc.), acquiring approximately 100-150 cells within 180×180 pixel field of view at ∼12.88 Hz frame rates. A Ti:Sapphire laser (Chameleon Ultra II, Coherent) operating at 920 nm provided excitation. We used a water-immersion objective (N16XLWD-PF - 16× Nikon CFI LWD Plan Fluorite, 0.80 NA, 3.0 mm WD) to focus the scanned laser illumination through the cranial window into the sample, and to collect the fluorescent signal from the GCaMP6f fluorophore expressed in pyramidal neurons. Photons were detected using an analog GaAsP photomultiplier tube (H7422P-40; Hamamatsu, Japan). Signal amplification utilized 2 µs pixel binning. Complete darkness during all experiments prevented light interference.

LabVIEW 8.0 (National Instruments) controlled imaging acquisition. To eliminate scan mirror auditory cues signaling trial transitions, mirrors remained active during inter-trial intervals. Arduino-generated TTL pulses synchronized imaging with behavioral events.

### CA1 neuronal activity analysis

#### Preprocessing raw images

Neuronal activity signals from the PMT were acquired using LabVIEW 8.0 in TIFF format. The MATLAB implementation of Suite2p (Pachitariu et al., 2016) was used to carry out semi-automated ROI detection including correction for frame-to-frame offsets due to sample motion.

Suite2p’s ROI identification algorithm clusters pixels that exhibit highly correlated intensity profiles within 10-15 µm size constraints, matching pyramidal neuron soma dimensions. Pixels within each ROI receive correlation-based weightings relative to centroid intensity, with cellular activity calculated as the weighted pixel sum. Suite2p also removed regions of overlap between cells, and excluded regions classified as neuropil. Manual ROI refinement utilized Suite2p functions. Custom MATLAB scripts (R2022b) were used to perform subsequent analyses.

#### dF/F Calculation

Trial-specific baseline fluorescence for each cell was determined as the 10th percentile of raw fluorescence. Frame-by-frame fluorescence values underwent baseline subtraction, normalization, and conversion to dF/F traces. These preprocessed datasets were stored as 3D matrices in .mat format, organized as cells × trials × frames.

#### dF/F Cleanup and Filtering

LED-flash artifacts from the conditioned stimulus in light-puff sessions contaminated all neuronal signals through objective capture. The CS-frame dF/F value was replaced with the median of the preceding five frames. The original recordings were cleaned up and truncated to 181 frames and aligned with respect to the CS frame. This effectively gave us the following time scale for each epoch: frames 1-58 defined as Pre-stim, starting with the CS frame at 59-60, variable trace frames as per protocol, and US (2 frames) together defined as the Stim period, and, finally, from frames ∼70 to 181 as Post-stim period.

#### Time Cell Detection

Time cells, defined as neurons exhibiting consistent trial-to-trial firing at fixed latencies relative to CS onset, were identified using custom C++ code (Ananthamurthy and Bhalla, 2023). This code is a fast implementation of two previously published algorithms for time-cell classification. The first is the ridge-to-background analysis (Modi et al., 2014), which was used for most of our analysis. The C++ code also obtains the temporal information metric (Mau et al., 2018b) to identify time cells.

For ridge-to-background analysis, each cell’s peak response time (PT) was extracted from the mean of the dF/F traces in the time interval from CS onset through 1.5 seconds post-US termination using alternate trials. The remaining trials computed ridge-to-background (R2B) ratio scores assessing reliability. The summed area encompassing the PT and two adjacent points in averaged traces was ratioed against area under all remaining points, defining the R2B ratio. Control measures randomly time-shifted traces before averaging, identifying independent PTs for each shift and computing corresponding R2B ratios. After 1,000 iterations per cell, averaged control values were obtained. R2B scores equaled the ratio of aligned-trace R2B ratios to randomly-shifted trace R2B ratios. Cells with R2B scores >2 received time cell classification.

### Behaviour Analysis

Camera-captured eyeblinks were saved as TIFF files. Custom MATLAB scripts identified and binarized eye regions to obtain a classification into eye vs non-eye regions. An ROI over the central region of the eye was analyzed for eyeblink status according to the method of Siegel et al. (2015). We estimated the Fraction Eye Closure (FEC) for each frame, defined as the ratio of eye-representing pixels to maximum fully-open-eye pixels for that session.

Per-trial baselines averaged FEC values across 500 ms pre-CS, minimizing false positives from partially-closed pre-CS eyes. Eyeblink responses were defined as FEC values exceeding 10% above baseline:

Baseline FEC = mean FEC (500 ms pre-CS) Eyeblink threshold = Baseline FEC + 0.10

FEC values exceeding the threshold between CS onset and US onset were classified as Conditioned Responses (CR). Trials exhibiting excessive pre-stimulus blinking (Fano factor >0.5) were excluded to prevent false positives.

### Statistical Analysis

Statistical analyses for Figures 3, 4E, 4F, 5 and 6 were performed using custom MATLAB scripts (R2022b, Academic Use). Statistical analyses for Figure 2 and 4A-D were performed in Python using the scipy.stats library. For Figure 4A-D we compared a 2-D array of normalized dF/F values between conditions. The arrays were obtained on a per-session basis, thus each pixel was averaged over all trials within a session. The rows were cells, and the columns were imaging frames. We took the vector of dF/F values for each cell within the selected time window (frame 59, CS to frame 98, CS+3s). We visually selected cells whose peak was within this time window. For each cell, we used Spearman’s test to compare the vectors between conditions, such as 250ms paired trials vs 550ms paired trials. We repeated this for each cell to obtain a vector of Spearman’s rho statistics. We aggregated the rho values for the population of cells and reported the median rho value, and then used the Wilcoxon signed-rank test on the distribution of rho values against zero to obtain a population significance level. In Figures 4A and 4B, we first eliminated the time-cells as they had a known correlation. Following this, the number of cells was still ∼5x the number of time cells, so we picked every 5th cell to perform the analysis with a comparable number of cells.

#### Population event rate and time cell tuning density (Figure 3)

Population event rate was defined as the fraction of cell-frames exceeding a per-cell, per-trial threshold of mean + 2 SD of the pre-CS baseline (frames 1–58). This binarisation was applied to the raw Gaussian-smoothed dF/F traces (dfbf_clean_noSigFilt) without additional filtering, and event rate was computed as the proportion of suprathreshold cell-frames within each temporal epoch.

Time cell tuning density was defined as the fraction of all recorded cells with a time-cell peak within a given epoch, divided by epoch duration (%/s). Peaks were identified as the argmax of the smoothed trial-averaged dF/F trace in the post-CS window (frames 59–181). Five epochs were defined relative to trial onset: Pre-CS (frames 1–58, 4.50 s), E1 (frames 59–71, 0–1 s post-CS), E2 (frames 72–97, 1–3 s), E3 (frames 98–136, 3–6 s), and E4 (frames 137–181, >6 s).

For within-protocol epoch comparisons (Figure 3Bii), the primary test was session-level bootstrapping (10,000 iterations, seed=42): sessions were resampled with replacement and tuning density recomputed each iteration, yielding 95% CIs and one-sided p-values for adjacent epoch decreases, with (count+1)/(N+1) continuity correction. Omnibus differences across epochs were assessed with the Friedman test on mouse-averaged values (n=4 mice), the appropriate repeated-measures non-parametric test given the same animals contributing to all epochs.

For cross-protocol comparisons (Figure 3Cii), the Friedman test was applied to mouse-averaged pooled post-CS density across all six protocols (n=4 mice). Because this omnibus test is underpowered at n=4 (minimum achievable Wilcoxon p=0.125), a supplementary bootstrap test directly compared all conditioning sessions pooled (n=111, across 5 protocols) against Extinction sessions (n=15): sessions were resampled with replacement (10,000 iterations, seed=42) and the difference in means recomputed each iteration; two-sided p was derived with (count+1)/(N+1) correction. Cohen’s d (pooled SD) was computed as an effect size measure. As sessions are not fully independent (the same 4 mice contributed across protocols), this session-level result is reported alongside the mouse-level omnibus.Tuning density computation and epoch definitions for Figure 3 follow the same procedure as described for Figures 5A–B below.

#### Tuning density and epoch distribution analyses (Figures 3B, 3C, 5A, 5B)

Tuning density was defined as the percentage of time cells with trial-averaged activity peaks within a given epoch, divided by epoch duration (%/s). Peak times were taken as the frame of maximum mean dF/F in the post-CS window (frames 59–181) for each time cell. Peaks were assigned to four epochs relative to CS onset: 0–1s, 1–3s, 3–6s, and >6s.

Non-uniform distribution of cells across epochs was tested using a chi-square goodness-of-fit test against the null of counts proportional to epoch duration (cell-level, descriptive). Monotonic trend across ordered epochs was assessed with the Cochran-Armitage trend test, using epoch midpoints (0.5, 2.0, 4.5, 8.0 s) as scores; one-sided (decreasing) for TC-250 and combined, two-sided for TC-550. Between-condition differences per epoch (TC-250 vs TC-550) were assessed with chi-square 2×2 tests (Fisher’s exact when expected counts < 5), also at the cell level.

For all inferential comparisons, session-level bootstrapping was the primary test. Sessions (n = 22) were resampled with replacement (10,000 iterations) and constituent cells pooled to recompute density per epoch, yielding bootstrap 95% CIs (2.5th–97.5th percentiles) and p-values. Adjacent-epoch decrease p-values used a one-sided test with (count + 1)/(N + 1) correction; between-condition p-values were two-sided. Because both the 250ms and 550ms peaks were drawn from the same resampled sessions in each iteration, within-session correlation between conditions was preserved. Overall peak time distributions (TC-250 vs TC-550, Figure 4F) were compared with two-sample Kolmogorov-Smirnov and Mann-Whitney U tests.

#### Epoch-specific scaling analysis (Figure 5C i-iii)

To compare peak times between conditions, cells significant as time cells in both the 250 ms and 550 ms conditions simultaneously were identified as common time cells (n = 30). For each common cell, the peak activity frame was determined from the trial-averaged dF/F trace for 250 ms trials and 550 ms trials separately.

Outliers were identified and excluded prior to regression using a shift-based MAD procedure applied within each epoch group (binned by 250 ms peak time). For each group, the shift was defined as the difference in peak frame between conditions (peak₅₅₀ − peak₂₅₀). The median absolute deviation (MAD) of shifts was computed and scaled by 1.4826 to yield a robust estimate of variability (σ); a minimum floor of 2 frames was applied to prevent over-exclusion in low-variability groups. Cells whose shift deviated more than 2.5σ from the group median shift were excluded as outliers (5 of 30 cells removed).

Linear regression of peak₅₅₀ on peak₂₅₀ was performed on the remaining valid cells, both for the overall population (restricted to cells with 250 ms peaks within 3 s of CS onset to avoid leverage from sparse late-peaking cells) and separately for cells originating in each epoch. Pearson correlation coefficients and regression slopes are reported with standard errors. To test whether timing was preserved (slope = 1), the fitted slope was tested against unity using a two-sided t-test (t = (slope − 1) / SE_slope, df = n − 2).

#### Tuning width analysis (Figure 5E)

FWHM was computed from each time cell’s smoothed trial-averaged dF/F trace in the post-CS window. Differences across epochs were assessed with the Kruskal-Wallis test; post-hoc pairwise comparisons used the Mann-Whitney U test across all six epoch pairs (exact p-values reported in Results).

## Acknowledgements

We acknowledge funding support from the Department of Biotechnology grant BT/PR12255/MED/122/8/2016, and NCBS-TIFR core funding from the Department of Atomic Energy, Government of India, Project Identification No. TRI 4006. We acknowledge NCBS campus facilities including Central Imaging and Flow Cytometry Facility, the Animal Care Resource Centre; and the mechanical workshop. We thank Soumya Bhattacharjee, Kambadur Ananthamurthy and Sriram Narayanan for help in imaging setup and analysis codes, and Bhanu Priya Somashekar and Anal Kumar for analysis.

## Author contributions

**Hrishikesh Nambisan:** Investigation, Software, Validation, Formal Analysis, Data Curation, Writing - Original Draft and Review and Editing, Visualisation.

**U.S.Bhalla:** Conceptualisation, Methodology, Software, Validation, Formal Analysis, Resources, Data Curation, Writing - Original Draft and Review and Editing, Visualisation, Supervision, Project Administration, Funding Acquisition.

## List of abbreviations

TEC: Trace Eyeblink Conditioning
CS: Conditioned Stimulus
US: Unconditioned Stimulus
CR: Conditioned Response
UR: Unconditioned Response
CA1: Cornu Ammonis area 1
PN: Pyramidal Neuron
BTSP: Behaviour Time Scale plasticity
FEC: Fraction Eye Closure
R2B: Ridge to Background

## Supplementary Material

**Supplementary Figure 1:**
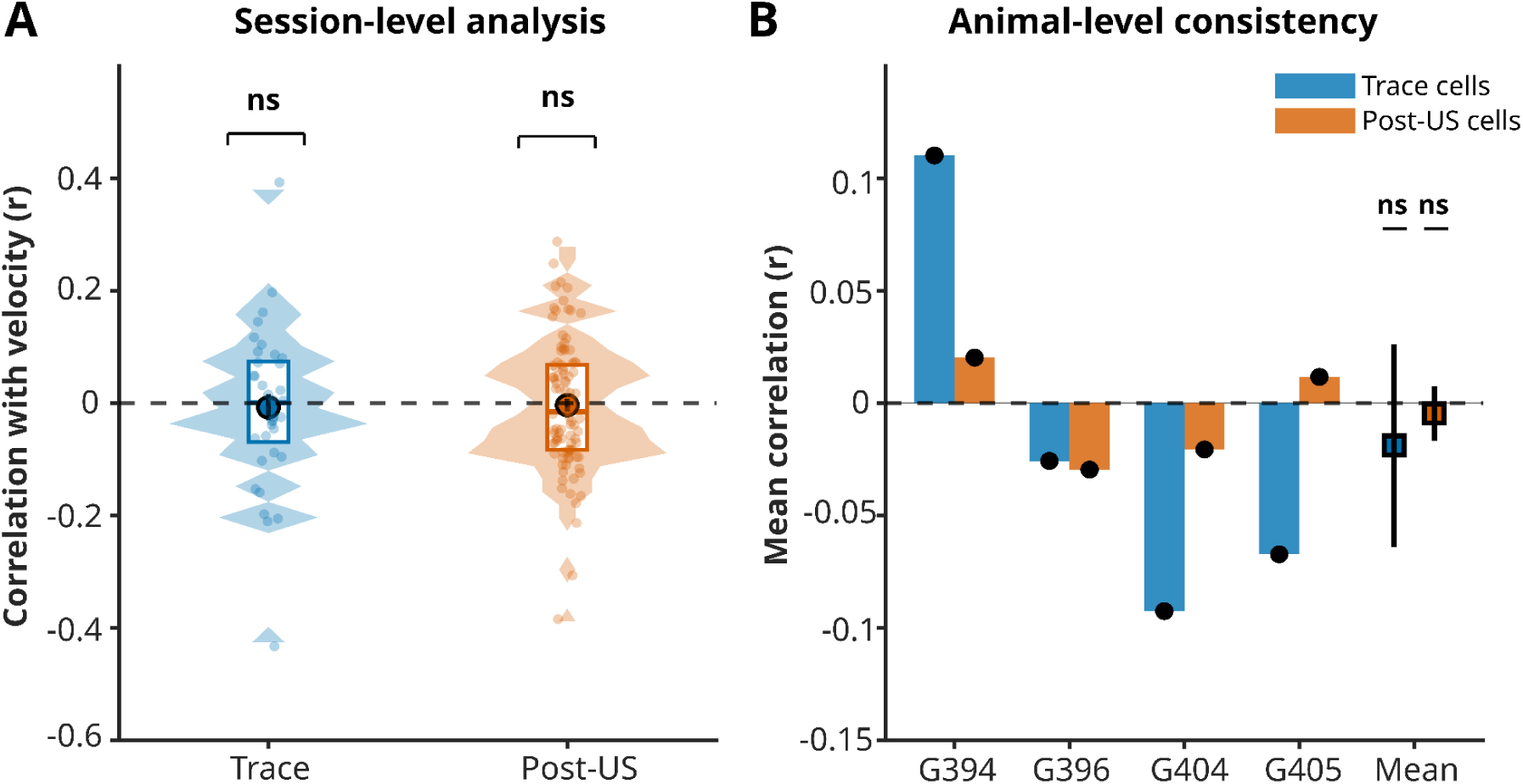
Treadmill movement does not correlate either with population or time-cell activity. **A:** The correlation of velocity with the population activity doesn’t change between Trace (r = -0.008±0.140) and post US period (r = -0.004±0.114). One-Sample Tests (against r=0): Trace vs 0: t-test p=0.7389 (ns); Post vs 0: t-test p=0.7360 (ns); Comparison (Post vs Trace): Paired sessions (N=36): Wilcoxon p=0.0000 ***. **B:** Animal level breakdown of A. One-Sample Tests (against r=0): Trace vs 0: t-test p=0.7041 (ns) Post vs 0: t-test p=0.7291 (ns) Comparison (Post vs Trace): Paired: Wilcoxon p=0.0000 ***.

**Supplementary Figure 2:**
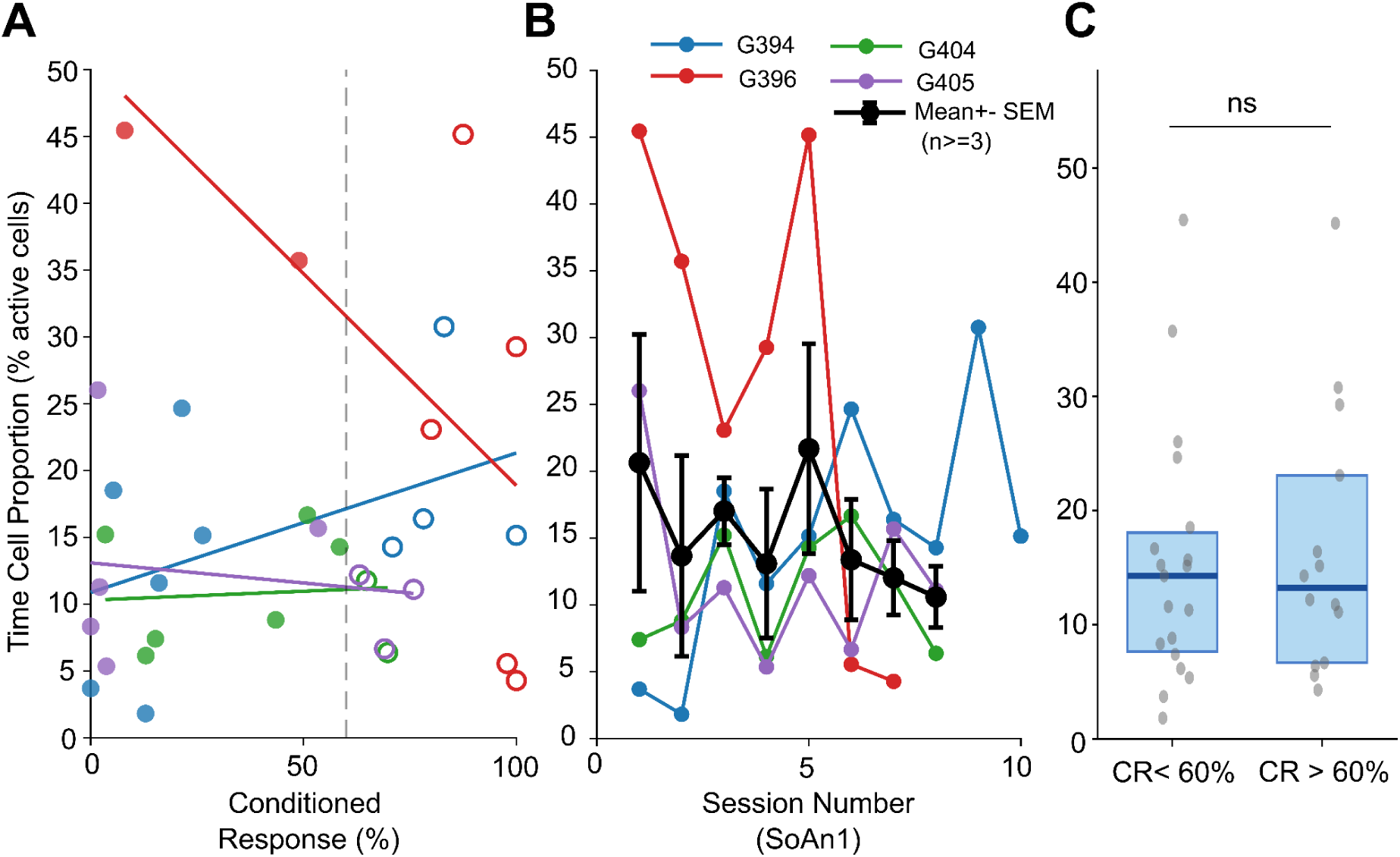
Time cell proportion is independent of learning for 250ms protocol. **A.** Scatter plot of conditioned response rate (CR%) versus time cell proportion (TC%; fraction of active CA1 cells meeting the R2B criterion) for all 250 ms trace sessions from four mice (G394: blue, G396: red, G404: green, G405: purple). Each point represents one session. Filled circles: pre-criterion sessions (CR < 60%); open circles: post-criterion sessions (CR ≥ 60%). Lines show within-animal linear regression fits. Dashed vertical line: 60% CR criterion. No significant relationship was found between CR% and TC proportion (within-animal Spearman correlations: G394 r = 0.46, p = 0.19; G396 r = −0.70, p = 0.09; G404 r = −0.02, p = 0.98; G405 r = −0.17, p = 0.70; group-level Fisher z-test: mean r = −0.14 ± 0.28 SEM, t(3) = −0.51, p = 0.65; linear mixed-effects model with mouse as random effect: β = −0.005 ± 0.053, t = −0.10, p = 0.92). **B.** Animal wise change in time cell proportions as sessions progress for 250 ms trace interval. **C** Comparison of TC proportion between pre-criterion (CR < 60%; n = 19 sessions, median = 14.3%, mean = 15.4 ± 9.5%) and post-criterion (CR ≥ 60%; n = 14 sessions, median = 13.2%, mean = 16.6 ± 10.4%) 250ms sessions. Box: median ± IQR; whiskers: 1.5 × IQR; points: individual sessions. No significant difference (Wilcoxon rank-sum, p = 0.87).

**Supplementary Figure 3:**
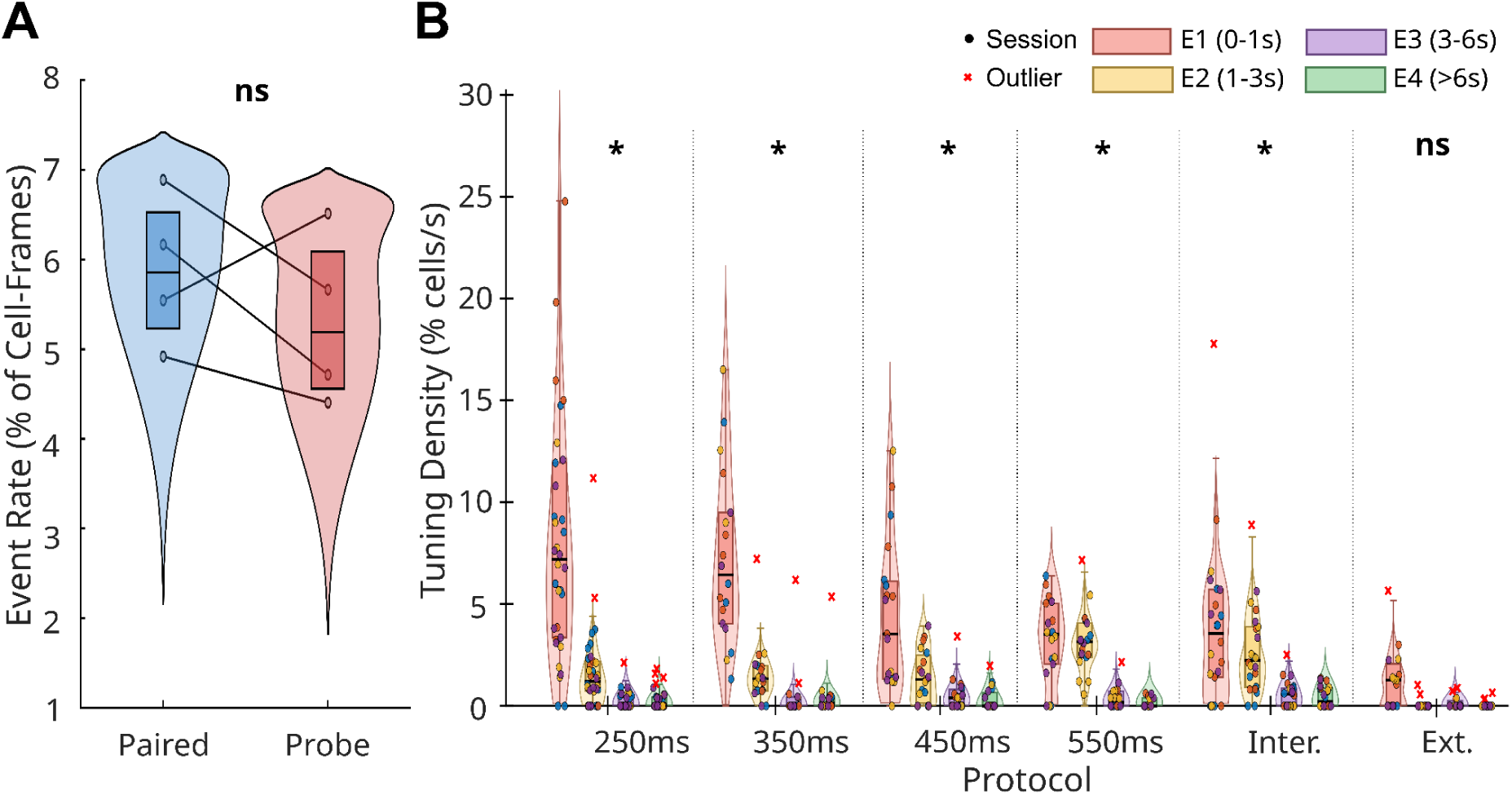
Time Cell Distributions **A:** Population event rate (fraction of suprathreshold cell-frames, 2 SD threshold) did not differ significantly between paired and probe trials (Wilcoxon signed-rank test, n=4 mice; p=0.375, ns). Lines connect matched mouse-averaged values across conditions.**B:** Time cell tuning density (%/s) for each combination of training protocol and post-CS temporal epoch (E1: 0–1 s; E2: 1–3 s; E3: 3–6 s; E4: >6 s). Each panel shows session-level distributions per protocol, with epochs colour-coded. Violin shapes show the kernel density estimate computed on non-outlier sessions; horizontal lines indicate the median; boxes indicate the IQR; whiskers extend to 1.5×IQR. Filled circles show individual sessions coloured by animal (n=4 mice). Red crosses indicate outlier sessions (>Q3+1.5×IQR), displayed at their true values and retained in all analyses.Within each protocol, tuning density differed significantly across epochs (Friedman test on mouse-averaged values, n=4 mice): 250 ms (χ²(4)=p=0.013), 350 ms (p=0.011), 450 ms (p=0.011), 550 ms (p=0.019), and IL (p=0.017). Extinction did not reach significance (p=0.072), consistent with the near-absent time cell activity in that protocol. In all conditioning protocols, density was highest in E1 and declined steeply across later epochs.Cross-protocol comparisons within each epoch revealed that differences were concentrated in the earliest post-CS window. E1 density differed significantly across protocols (Friedman χ²(5)=14.86, p=0.011); post-hoc Bonferroni comparisons identified 250 ms (p=0.010) and 350 ms (p=0.038) as significantly exceeding Extinction. E2 density also differed across protocols (Friedman χ²(5)=11.43, p=0.044), with 550 ms exceeding Extinction after Bonferroni correction (p=0.020). E3 and E4 showed no significant cross-protocol differences (p=0.186 and p=0.167, respectively).

**Supplementary Figure 4:**
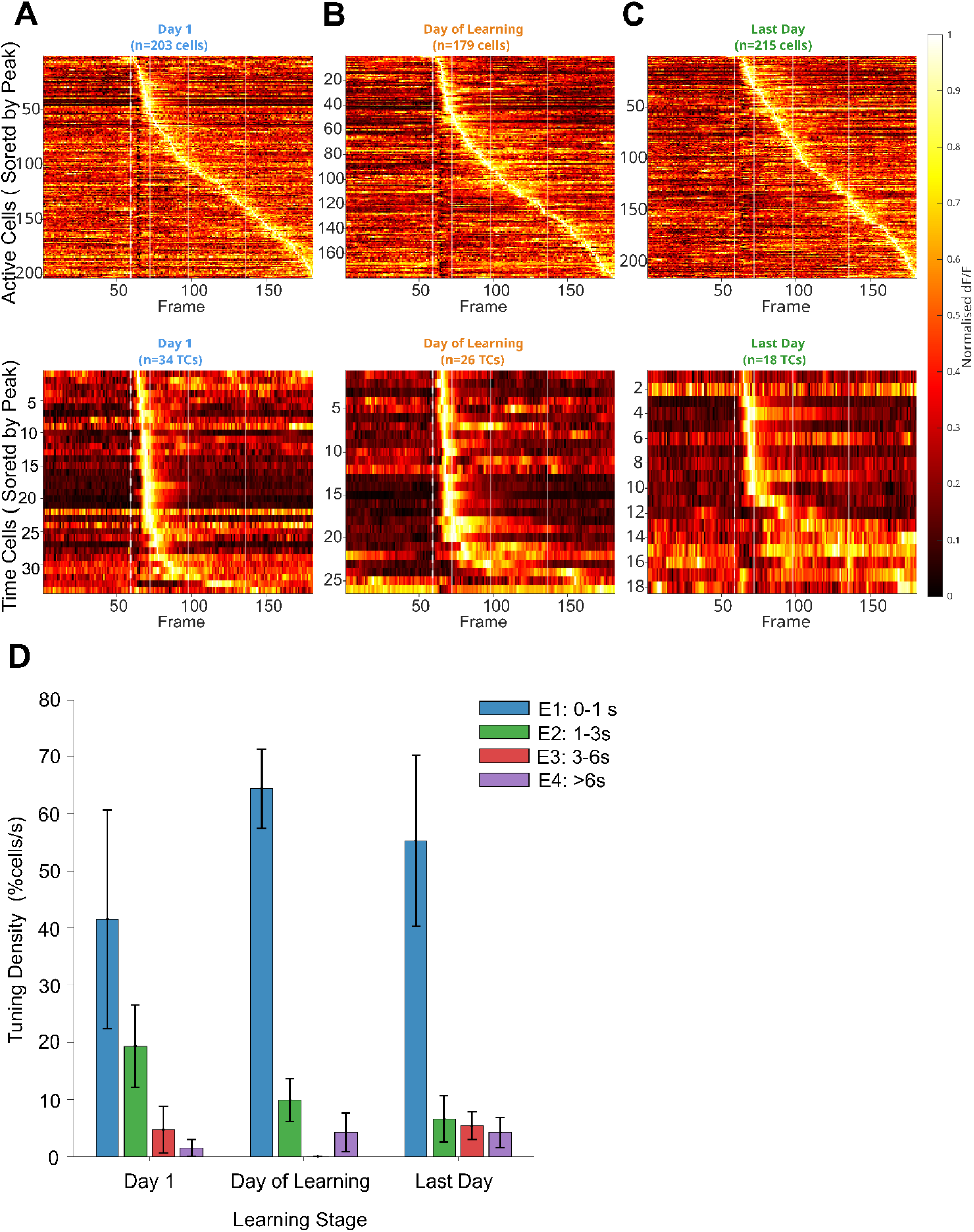
Time cell tuning density distribution across epochs remains consistent across learning. **(A–C)** Trial-averaged population activity heatmaps for active cells (top row) and time cells (R2B criterion, bottom row) pooled across all four mice, sorted by peak frame (post-CS window), for three SoAn1 learning stages: Day 1 (first session for each animal; n = 35 time cells from 203 active cells), Day of Learning (first session where CR ≥ 60% per animal; n = 26 time cells from 179 active cells), and Last Day (final SoAn1 session; n = 21 time cells from 215 active cells). Target sessions per animal: Day 1 — G394 Day 2 (CR = 0%), G396 Day 2 (CR = 8%), G404 Day 2 (CR = 15%), G405 Day 3 (CR = 2%); Day of Learning — G394 Day 8 (CR = 78%), G396 Day 4 (CR = 80%), G404 Day 8 (CR = 65%), G405 Day 7 (CR = 63%). Colour scale: normalised dF/F, 0–1 per cell. Dashed white vertical line: CS onset (frame 59). Sequential structure spanning the full post-CS window is visible across all stages. Time cell count decreases modestly from Day 1 to Last Day (35 → 26 → 21), consistent with gradual turnover rather than learning-dependent recruitment. **(D)** Epoch-specific time cell tuning density (% time cells per second) across the four post-CS epochs (E1: 0–1 s, E2: 1–3 s, E3: 3–6 s, E4: >6 s) for each learning stage. For each session, time cells were identified by TcPy R2B sigBootstrap criterion; peak frames were computed from Gaussian-smoothed (window = 5 frames) trial-averaged dF/F traces over the full post-CS window (frames 59–181). Density = (fraction of time cells peaking in epoch) / epoch duration (s). Bars show mean ± SEM across four sessions (one per mouse per stage). E1 (0–1 s) consistently dominates across all three stages (Day 1: mean = 41.5 ± 19.1 %TCs/s; Day of Learning: 64.4 ± 6.9 %TCs/s; Last Day: 55.3 ± 15.0 %TCs/s). No significant cross-stage differences were detected for any epoch (Wilcoxon rank-sum, all p > 0.14). Within-stage Friedman tests across epochs were significant at Day of Learning (χ²(3) = 9.81, p = 0.020) and Last Day (χ²(3) = 8.33, p = 0.040), and showed a trend at Day 1 (χ²(3) = 7.05, p = 0.070), driven by E1 dominance in all stages; however, no individual post-hoc pairwise comparison survived Dunn-Šidák correction (adjusted α = 0.0085). Note: statistical power is limited (n = 4 sessions per stage); cross-stage comparisons should be interpreted with caution.All analyses use paired trials. Cells normalised [0, 1] per cell prior to pooling. Frame rate: 12.88 Hz; CS onset: frame 59 of 181 total frames. Epoch boundaries: E1 frames 59–71, E2 frames 72–97, E3 frames 98–135, E4 frames 136–181.

## Notes

### Competing Interest Statement

The authors have declared no competing interest.

## References

Ahmed, M. S., Priestley, J. B., Castro, A., Stefanini, F., Solis Canales, A. S., Balough, E. M., Lavoie, E., Mazzucato, L., Fusi, S., & Losonczy, A. (2020). Hippocampal Network Reorganization Underlies the Formation of a Temporal Association Memory. Neuron, 107(2), 283–291.e6. 10.1016/j.neuron.2020.04.013

Ananthamurthy, K. G., & Bhalla, U. S. (2023). Synthetic Data Resource and Benchmarks for Time Cell Analysis and Detection Algorithms. Eneuro, 10(3), ENEURO.0007–22.2023. 10.1523/ENEURO.0007-22.2023

Azizi, A. H., Wiskott, L., & Cheng, S. (2013). A computational model for preplay in the hippocampus. Frontiers in Computational Neuroscience, 7. 10.3389/fncom.2013.00161

Berger, T. W., Alger, B., & Thompson, R. F. (1976). Neuronal Substrate of Classical Conditioning in the Hippocampus. Science, 192(4238), 483–485. 10.1126/science.1257783

Berger, T. W., Rinaldi, P. C., Weisz, D. J., & Thompson, R. F. (1983). Single-unit analysis of different hippocampal cell types during classical conditioning of rabbit nictitating membrane response. Journal of Neurophysiology, 50(5), 1197–1219. 10.1152/jn.1983.50.5.1197

Bhalla, U. S. (2002). Mechanisms for Temporal Tuning and Filtering by Postsynaptic Signaling Pathways. Biophysical Journal, 83(2), 740–752. 10.1016/S0006-3495(02)75205-3

Bhattacharjee, S., Nambisan, H., & Bhalla, U. S. (2024). Time cells emerge early in learning and encode stimulus modality past task requirements. bioRxiv, 2024.07.28.605458. 10.1101/2024.07.28.605458

Bigus, E. R., Lee, H.-W., Bowler, J. C., Shi, J., & Heys, J. G. (2024). Medial entorhinal cortex plays a specialized role in learning of flexible, context-dependent interval timing behavior (p. 2023.01.18.524598). bioRxiv. 10.1101/2023.01.18.524598

Bittner, K. C., Grienberger, C., Vaidya, S. P., Milstein, A. D., Macklin, J. J., Suh, J., Tonegawa, S., & Magee, J. C. (2015). Conjunctive input processing drives feature selectivity in hippocampal CA1 neurons. Nature Neuroscience, 18(8), 1133–1142. 10.1038/nn.4062

Bittner, K. C., Milstein, A. D., Grienberger, C., Romani, S., & Magee, J. C. (2017). Behavioral time scale synaptic plasticity underlies CA1 place fields. Science, 357(6355), 1033–1036. 10.1126/science.aan3846

Bostock, E., Muller, R. U., & Kubie, J. L. (1991). Experience-dependent modifications of hippocampal place cell firing. Hippocampus, 1(2), 193–205. 10.1002/hipo.450010207

Bouton, M. E. (2004). Context and Behavioral Processes in Extinction. Learning & Memory, 11(5), 485–494. 10.1101/lm.78804

Bouton, M. E., & Bolles, R. C. (1979). Contextual control of the extinction of conditioned fear. Learning and Motivation, 10(4), 445–466. 10.1016/0023-9690(79)90057-2

Buonomano, D. V., & Laje, R. (2010). Population clocks: Motor timing with neural dynamics. Trends in Cognitive Sciences, 14(12), 520–527. 10.1016/j.tics.2010.09.002

Buonomano, D. V., & Maass, W. (2009). State-dependent computations: Spatiotemporal processing in cortical networks. Nature Reviews Neuroscience, 10(2), 113–125. 10.1038/nrn2558

Cao, R., Bladon, J. H., Charczynski, S. J., Hasselmo, M. E., & Howard, M. W. (2022). Internally generated time in the rodent hippocampus is logarithmically compressed. eLife, 11, e75353. 10.7554/eLife.75353

Clark, R. E., Manns, J. R., & Squire, L. R. (2002). Classical conditioning, awareness, and brain systems. Trends in Cognitive Sciences, 6(12), 524–531. 10.1016/S1364-6613(02)02041-7

Dias, M., Ferreira, R., & Remondes, M. (2021). Medial Entorhinal Cortex Excitatory Neurons Are Necessary for Accurate Timing. The Journal of Neuroscience, 41(48), 9932–9943. 10.1523/JNEUROSCI.0750-21.2021

Diesmann, M., Gewaltig, M.-O., & Aertsen, A. (1999). Stable propagation of synchronous spiking in cortical neural networks. Nature, 402(6761), 529–533. 10.1038/990101

Dragoi, G., & Buzsáki, G. (2006). Temporal Encoding of Place Sequences by Hippocampal Cell Assemblies. Neuron, 50(1), 145–157. 10.1016/j.neuron.2006.02.023

Dragoi, G., & Tonegawa, S. (2011). Preplay of future place cell sequences by hippocampal cellular assemblies. Nature, 469(7330), 397–401. 10.1038/nature09633

Eichenbaum, H. (2014). Time cells in the hippocampus: A new dimension for mapping memories. Nature Reviews Neuroscience, 15(11), 732–744. 10.1038/nrn3827

Eichenbaum, H. (2017). On the Integration of Space, Time, and Memory. Neuron, 95(5), 1007–1018. 10.1016/j.neuron.2017.06.036

Frank, L. M., Brown, E. N., & Wilson, M. (2000). Trajectory Encoding in the Hippocampus and Entorhinal Cortex. Neuron, 27(1), 169–178. 10.1016/S0896-6273(00)00018-0

Frank, L. M., Stanley, G. B., & Brown, E. N. (2004). Hippocampal Plasticity across Multiple Days of Exposure to Novel Environments. Journal of Neuroscience, 24(35), 7681–7689. 10.1523/JNEUROSCI.1958-04.2004

Gill, P. R., Mizumori, S. J. Y., & Smith, D. M. (2011). Hippocampal episode fields develop with learning. Hippocampus, 21(11), 1240–1249. 10.1002/hipo.20832

Guger, C., Gener, T., Pennartz, C. M. A., Brotons-Mas, J. R., Edlinger, G., Bermúdez I Badia, S., Verschure, P., Schaffelhofer, S., & Sanchez-Vives, M. V. (2011). Real-time position reconstruction with hippocampal place cells. Frontiers in Neuroscience, 5, 85. 10.3389/fnins.2011.00085

Hasselmo, M. E., & Stern, C. E. (2015). Current questions on space and time encoding. Hippocampus, 25(6), 744–752. 10.1002/hipo.22454

Hill, A. J. (1978). First occurrence of hippocampal spatial firing in a new environment. Experimental Neurology, 62(2), 282–297. 10.1016/0014-4886(78)90058-4

Howard, M. W., & Eichenbaum, H. (2013). The hippocampus, time, and memory across scales. Journal of Experimental Psychology. General, 142(4), 1211–1230. 10.1037/a0033621

Howard, M. W., & Eichenbaum, H. (2015). Time and space in the hippocampus. Brain Research, 1621, 345–354. 10.1016/j.brainres.2014.10.069

Howard, M. W., MacDonald, C. J., Tiganj, Z., Shankar, K. H., Du, Q., Hasselmo, M. E., & Eichenbaum, H. (2014). A Unified Mathematical Framework for Coding Time, Space, and Sequences in the Hippocampal Region. The Journal of Neuroscience, 34(13), 4692–4707. 10.1523/JNEUROSCI.5808-12.2014

Kishimoto, Y., Kawahara, S., Mori, H., Mishina, M., & Kirino, Y. (2001). Long-trace interval eyeblink conditioning is impaired in mutant mice lacking the NMDA receptor subunit ε1. European Journal of Neuroscience, 13(6), 1221–1227. 10.1046/j.0953-816x.2001.01486.x

Komorowski, R. W., Manns, J. R., & Eichenbaum, H. (2009). Robust Conjunctive Item–Place Coding by Hippocampal Neurons Parallels Learning What Happens Where. The Journal of Neuroscience, 29(31), 9918–9929. 10.1523/JNEUROSCI.1378-09.2009

Kraus, B. J., Robinson, R. J., White, J. A., Eichenbaum, H., & Hasselmo, M. E. (2013). Hippocampal “Time Cells”: Time versus Path Integration. Neuron, 78(6), 1090–1101. 10.1016/j.neuron.2013.04.015

Kumar, A., Rotter, S., & Aertsen, A. (2008). Conditions for propagating synchronous spiking and asynchronous firing rates in a cortical network model. The Journal of Neuroscience, 28(20), 5268–5280. 10.1523/jneurosci.2542-07.2008

Kumar, A., Rotter, S., & Aertsen, A. (2010). Spiking activity propagation in neuronal networks: Reconciling different perspectives on neural coding. Nature Reviews Neuroscience, 11(9), 615–627. 10.1038/nrn2886

Laje, R., & Buonomano, D. V. (2013). Robust timing and motor patterns by taming chaos in recurrent neural networks. Nature Neuroscience, 16(7), 925–933. 10.1038/nn.3405

Lee, I., Griffin, A. L., Zilli, E. A., Eichenbaum, H., & Hasselmo, M. E. (2006). Gradual Translocation of Spatial Correlates of Neuronal Firing in the Hippocampus toward Prospective Reward Locations. Neuron, 51(5), 639–650. 10.1016/j.neuron.2006.06.033

Leutgeb, S., Leutgeb, J. K., Barnes, C. A., Moser, E. I., McNaughton, B. L., & Moser, M.-B. (2005). Independent Codes for Spatial and Episodic Memory in Hippocampal Neuronal Ensembles. Science, 309(5734), 619–623. 10.1126/science.1114037

Lever, C., Wills, T., Cacucci, F., Burgess, N., & O’Keefe, J. (2002). Long-term plasticity in hippocampal place-cell representation of environmental geometry. Nature, 416(6876), 90–94. 10.1038/416090a

Lt, T., & Pj, B. (1990). Long-term stability of the place-field activity of single units recorded from the dorsal hippocampus of freely behaving rats. Brain Research, 509(2). 10.1016/0006-8993(90)90555-p

MacDonald, C. J., Carrow, S., Place, R., & Eichenbaum, H. (2013). Distinct Hippocampal Time Cell Sequences Represent Odor Memories in Immobilized Rats. The Journal of Neuroscience, 33(36), 14607–14616. 10.1523/JNEUROSCI.1537-13.2013

MacDonald, C. J., Lepage, K. Q., Eden, U. T., & Eichenbaum, H. (2011). Hippocampal “Time Cells” Bridge the Gap in Memory for Discontiguous Events. Neuron, 71(4), 737–749. 10.1016/j.neuron.2011.07.012

Mankin, E. A., Sparks, F. T., Slayyeh, B., Sutherland, R. J., Leutgeb, S., & Leutgeb, J. K. (2012). Neuronal code for extended time in the hippocampus. Proceedings of the National Academy of Sciences, 109(47), 19462–19467. 10.1073/pnas.1214107109

Masuda, A., Sano, C., Zhang, Q., Goto, H., McHugh, T. J., Fujisawa, S., & Itohara, S. (2020). The hippocampus encodes delay and value information during delay-discounting decision making. eLife, 9, e52466. 10.7554/eLife.52466

Mau, W., Sullivan, D. W., Kinsky, N. R., Hasselmo, M. E., Howard, M. W., & Eichenbaum, H. (2018). The Same Hippocampal CA1 Population Simultaneously Codes Temporal Information over Multiple Timescales. Current Biology: CB, 28(10), 1499–1508.e4. 10.1016/j.cub.2018.03.051

McEchron, M. D., Bouwmeester, H., Tseng, W., Weiss, C., & Disterhoft, J. F. (1999). Hippocampectomy disrupts auditory trace fear conditioning and contextual fear conditioning in the rat. Hippocampus, 8(6), 638–646. 10.1002/(SICI)1098-1063(1998)8:6<638::AID-HIPO6>3.0.CO;2-Q

McEchron, M. D., & Disterhoft, J. F. (1997). Sequence of Single Neuron Changes in CA1 Hippocampus of Rabbits During Acquisition of Trace Eyeblink Conditioned Responses. Journal of Neurophysiology, 78(2), 1030–1044. 10.1152/jn.1997.78.2.1030

Modi, M. N., Dhawale, A. K., & Bhalla, U. S. (2014). CA1 cell activity sequences emerge after reorganization of network correlation structure during associative learning. eLife, 3, e01982. 10.7554/eLife.01982

Moyer, J. R., Deyo, R. A., & Disterhoft, J. F. (1990). Hippocampectomy disrupts trace eye-blink conditioning in rabbits. Behavioral Neuroscience, 104(2), 243–252. 10.1037/0735-7044.104.2.243

Muller, R., & Kubie, J. (1987). The effects of changes in the environment on the spatial firing of hippocampal complex-spike cells. The Journal of Neuroscience, 7(7), 1951–1968. 10.1523/JNEUROSCI.07-07-01951.1987

Myers, K. M., & Davis, M. (2007). Mechanisms of fear extinction. Molecular Psychiatry, 12(2), 120–150. 10.1038/sj.mp.4001939

O’Keefe, J., & Burgess, N. (1996). Geometric determinants of the place fields of hippocampal neurons. Nature, 381(6581), 425–428. 10.1038/381425a0

O’Keefe, J., & Recce, M. L. (1993). Phase relationship between hippocampal place units and the EEG theta rhythm. Hippocampus, 3(3), 317–330. 10.1002/hipo.450030307

Pastalkova, E., Itskov, V., Amarasingham, A., & Buzsáki, G. (2008). Internally Generated Cell Assembly Sequences in the Rat Hippocampus. Science, 321(5894), 1322–1327. 10.1126/science.1159775

Rahman, M. M., Shukla, A., & Chattarji, S. (2018). Extinction recall of fear memories formed before stress is not affected despite higher theta activity in the amygdala. eLife, 7, e35450. 10.7554/eLife.35450

Rajan, K., Harvey, C. D., & Tank, D. W. (2016). Recurrent Network Models of Sequence Generation and Memory. Neuron, 90(1), 128–142. 10.1016/j.neuron.2016.02.009

Rescorla, R. A. (2004). Spontaneous Recovery. Learning & Memory, 11(5), 501–509. 10.1101/lm.77504

Sabariego, M., Schönwald, A., Boublil, B. L., Zimmerman, D. T., Ahmadi, S., Gonzalez, N., Leibold, C., Clark, R. E., Leutgeb, J. K., & Leutgeb, S. (2019). Time Cells in the Hippocampus Are Neither Dependent on Medial Entorhinal Cortex Inputs nor Necessary for Spatial Working Memory. Neuron, 102(6), 1235–1248.e5. 10.1016/j.neuron.2019.04.005

Salz, D. M., Tiganj, Z., Khasnabish, S., Kohley, A., Sheehan, D., Howard, M. W., & Eichenbaum, H. (2016). Time Cells in Hippocampal Area CA3. Journal of Neuroscience, 36(28), 7476–7484. 10.1523/JNEUROSCI.0087-16.2016

Siegel, J. J., Taylor, W., Gray, R., Kalmbach, B., Zemelman, B. V., Desai, N. S., Johnston, D., & Chitwood, R. A. (2015). Trace Eyeblink Conditioning in Mice Is Dependent upon the Dorsal Medial Prefrontal Cortex, Cerebellum, and Amygdala: Behavioral Characterization and Functional Circuitry,,. eNeuro, 2(4), ENEURO.0051–14.2015. 10.1523/ENEURO.0051-14.2015

Skaggs, W. E., McNaughton, B. L., Wilson, M. A., & Barnes, C. A. (1996). Theta phase precession in hippocampal neuronal populations and the compression of temporal sequences. Hippocampus, 6(2), 149–172. 10.1002/(SICI)1098-1063(1996)6:2<149::AID-HIPO6>3.0.CO;2-K

Solomon, P. R., Vander Schaaf, E. R., Thompson, R. F., & Weisz, D. J. (1986). Hippocampus and trace conditioning of the rabbit’s classically conditioned nictitating membrane response. Behavioral Neuroscience, 100(5), 729–744. 10.1037/0735-7044.100.5.729

Taxidis, J., Pnevmatikakis, E. A., Dorian, C. C., Mylavarapu, A. L., Arora, J. S., Samadian, K. D., Hoffberg, E. A., & Golshani, P. (2020). Differential Emergence and Stability of Sensory and Temporal Representations in Context-Specific Hippocampal Sequences. Neuron, 108(5), 984–998.e9. 10.1016/j.neuron.2020.08.028

Tseng, W., Guan, R., Disterhoft, J. f., & Weiss, C. (2004). Trace eyeblink conditioning is hippocampally dependent in mice. Hippocampus, 14(1), 58–65. 10.1002/hipo.10157

Tsodyks, M. V., & Markram, H. (1997). The neural code between neocortical pyramidal neurons depends on neurotransmitter release probability. Proceedings of the National Academy of Sciences, 94(2), 719–723. 10.1073/pnas.94.2.719

Umbach, G., Kantak, P., Jacobs, J., Kahana, M., Pfeiffer, B. E., Sperling, M., & Lega, B. (2020). Time cells in the human hippocampus and entorhinal cortex support episodic memory. Proceedings of the National Academy of Sciences, 117(45), 28463–28474. 10.1073/pnas.2013250117

Vaz, A. P., Wittig, J. H., Inati, S. K., & Zaghloul, K. A. (2023). Backbone spiking sequence as a basis for preplay, replay, and default states in human cortex. Nature Communications, 14(1), 4723. 10.1038/s41467-023-40440-5

Villette, V., Malvache, A., Tressard, T., Dupuy, N., & Cossart, R. (2015). Internally Recurring Hippocampal Sequences as a Population Template of Spatiotemporal Information. Neuron, 88(2), 357–366. 10.1016/j.neuron.2015.09.052

Weiss, C., Bouwmeester, H., Power, J. M., & Disterhoft, J. F. (1999). Hippocampal lesions prevent trace eyeblink conditioning in the freely moving rat. Behavioural Brain Research, 99(2), 123–132. 10.1016/S0166-4328(98)00096-5

Wilson, M. A., & McNaughton, B. L. (1993). Dynamics of the Hippocampal Ensemble Code for Space. Science, 261(5124), 1055–1058. 10.1126/science.8351520

Wood, E. R., Dudchenko, P. A., Robitsek, R. J., & Eichenbaum, H. (2000). Hippocampal Neurons Encode Information about Different Types of Memory Episodes Occurring in the Same Location. Neuron, 27(3), 623–633. 10.1016/S0896-6273(00)00071-4

Yuan, L., Figueroa, J. F., Khan, A., Narayan, G., Leutgeb, J. K., & Leutgeb, S. (2025). Time cell sequences during delay intervals are not dependent on brain state and do not support hippocampus-dependent working memory. Nature Communications, 16(1), 7470. 10.1038/s41467-025-62498-z

Ziv, Y., Burns, L. D., Cocker, E. D., Hamel, E. O., Ghosh, K. K., Kitch, L. J., Gamal, A. E., & Schnitzer, M. J. (2013). Long-term dynamics of CA1 hippocampal place codes. Nature Neuroscience, 16(3), 264–266. 10.1038/nn.3329

