## Supplementery Figure for "Time cells differentially populate trace and post-trace epochs but do not remap for different trace intervals"

### Supplementary Material

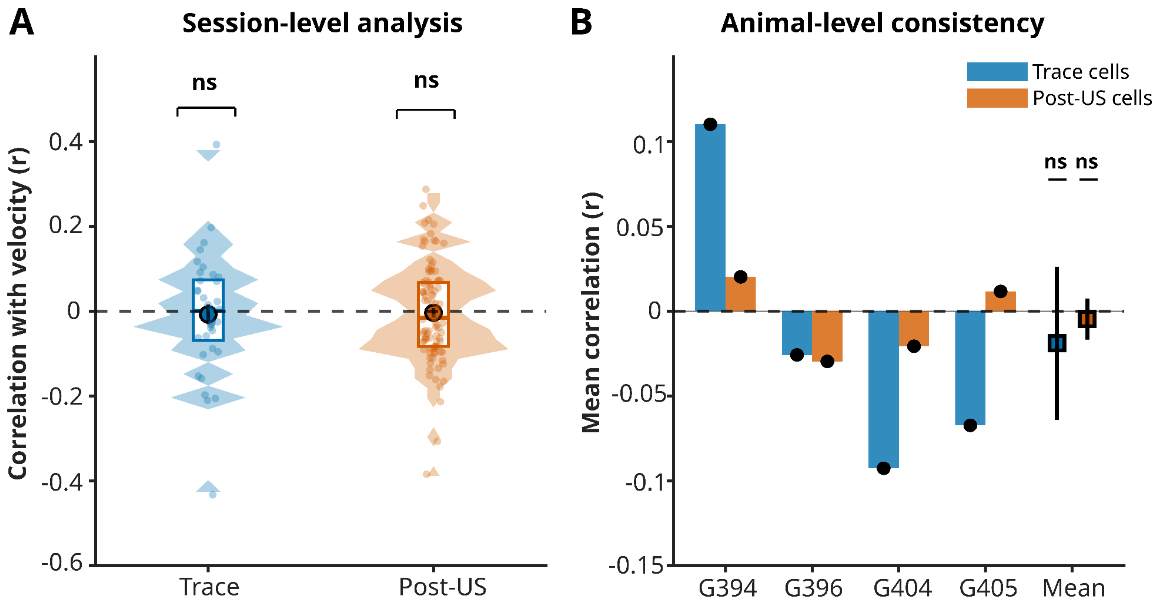


**Supplementary Figure 1**: Treadmill movement does not correlate either with population or time-cell activity. **A:** The correlation of velocity with the population activity doesn't change between Trace ( r = -0.008±0.140) and post US period (r = -0.004±0.114). One-Sample Tests (against r=0): Trace vs 0: t-test p=0.7389 (ns); Post vs 0: t-test p=0.7360 (ns); Comparison (Post vs Trace): Paired sessions (N=36): Wilcoxon p=0.0000 ***. **B:** Animal level breakdown of A. One-Sample Tests (against r=0): Trace vs 0: t-test p=0.7041 (ns) Post vs 0: t-test p=0.7291 (ns) Comparison (Post vs Trace): Paired: Wilcoxon p=0.0000 ***.

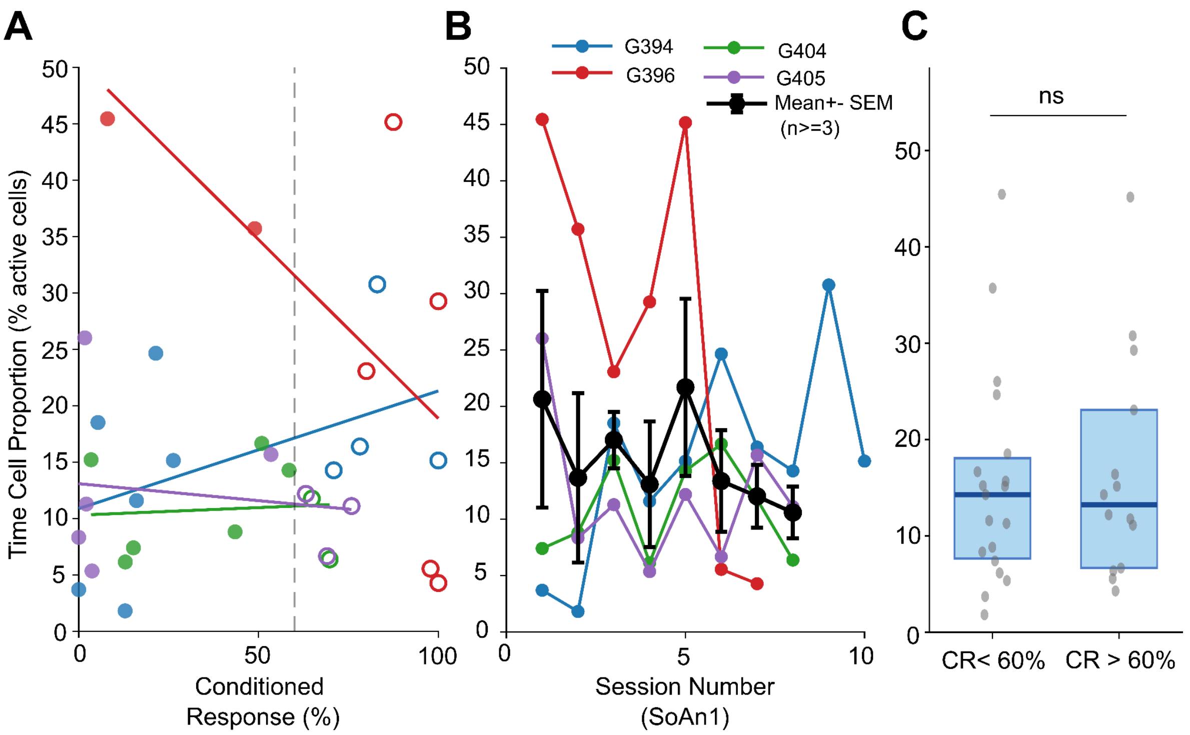


**Supplementary Figure 2:** Time cell proportion is independent of learning for 250ms protocol. **A.** Scatter plot of conditioned response rate (CR%) versus time cell proportion (TC%; fraction of active CA1 cells meeting the R2B criterion) for all 250 ms trace sessions from four mice (G394: blue, G396: red, G404: green, G405: purple). Each point represents one session. Filled circles: pre-criterion sessions (CR < 60%); open circles: post-criterion sessions (CR ≥ 60%). Lines show within-animal linear regression fits. Dashed vertical line: 60% CR criterion. No significant relationship was found between CR% and TC proportion (within-animal Spearman correlations: G394 $r = 0.46$, $p = 0.19$; G396 $r = -0.70$, $p = 0.09$; G404 $r = -0.02$, $p = 0.98$; G405 $r = -0.17$, $p = 0.70$; group-level Fisher z-test: mean $r = -0.14 \pm 0.28$ SEM, $t(3) = -0.51$, $p = 0.65$; linear mixed-effects model with mouse as random effect: $\beta = -0.005 \pm 0.053$, $t = -0.10$, $p = 0.92$). **B.** Animal wise change in time cell proportions as sessions progress for 250 ms trace interval. **C** Comparison of TC proportion between pre-criterion (CR < 60%; n = 19 sessions, median = 14.3%, mean = 15.4 ± 9.5%) and post-criterion (CR ≥ 60%; n = 14 sessions, median = 13.2%, mean = 16.6 ± 10.4%) 250ms sessions. Box: median ± IQR; whiskers: 1.5 × IQR; points: individual sessions. No significant difference (Wilcoxon rank-sum, $p = 0.87$).

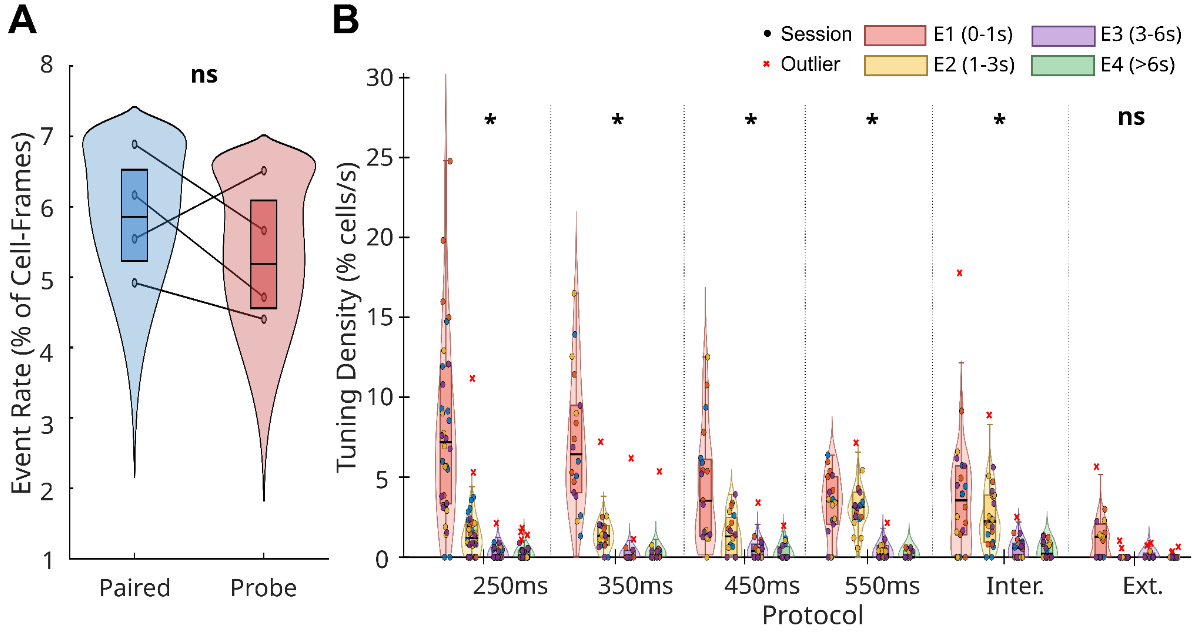


**Supplementary Figure 3:** Time Cell Distributions **A:** Population event rate (fraction of suprathreshold cell-frames, 2 SD threshold) did not differ significantly between paired and probe trials (Wilcoxon signed-rank test, n=4 mice; p=0.375, ns). Lines connect matched mouse-averaged values across conditions.**B:** Time cell tuning density (%/s) for each combination of training protocol and post-CS temporal epoch (E1: 0–1 s; E2: 1–3 s; E3: 3–6 s; E4: >6 s). Each panel shows session-level distributions per protocol, with epochs colour-coded. Violin shapes show the kernel density estimate computed on non-outlier sessions; horizontal lines indicate the median; boxes indicate the IQR; whiskers extend to 1.5×IQR. Filled circles show individual sessions coloured by animal (n=4 mice). Red crosses indicate outlier sessions (>Q3+1.5×IQR), displayed at their true values and retained in all analyses.Within each protocol, tuning density differed significantly across epochs (Friedman test on mouse-averaged values, n=4 mice): 250 ms ($\chi^2(4)$=p=0.013), 350 ms (p=0.011), 450 ms (p=0.011), 550 ms (p=0.019), and IL (p=0.017). Extinction did not reach significance (p=0.072), consistent with the near-absent time cell activity in that protocol. In all conditioning protocols, density was highest in E1 and declined steeply across later epochs.Cross-protocol comparisons within each epoch revealed that differences were concentrated in the earliest post-CS window. E1 density differed significantly across protocols (Friedman $\chi^2(5)$=14.86, p=0.011); post-hoc Bonferroni comparisons identified 250 ms (p=0.010) and 350 ms (p=0.038) as significantly exceeding Extinction. E2 density also differed across protocols (Friedman $\chi^2(5)$=11.43, p=0.044), with 550 ms exceeding Extinction after Bonferroni correction (p=0.020). E3 and E4 showed no significant cross-protocol differences (p=0.186 and p=0.167, respectively).

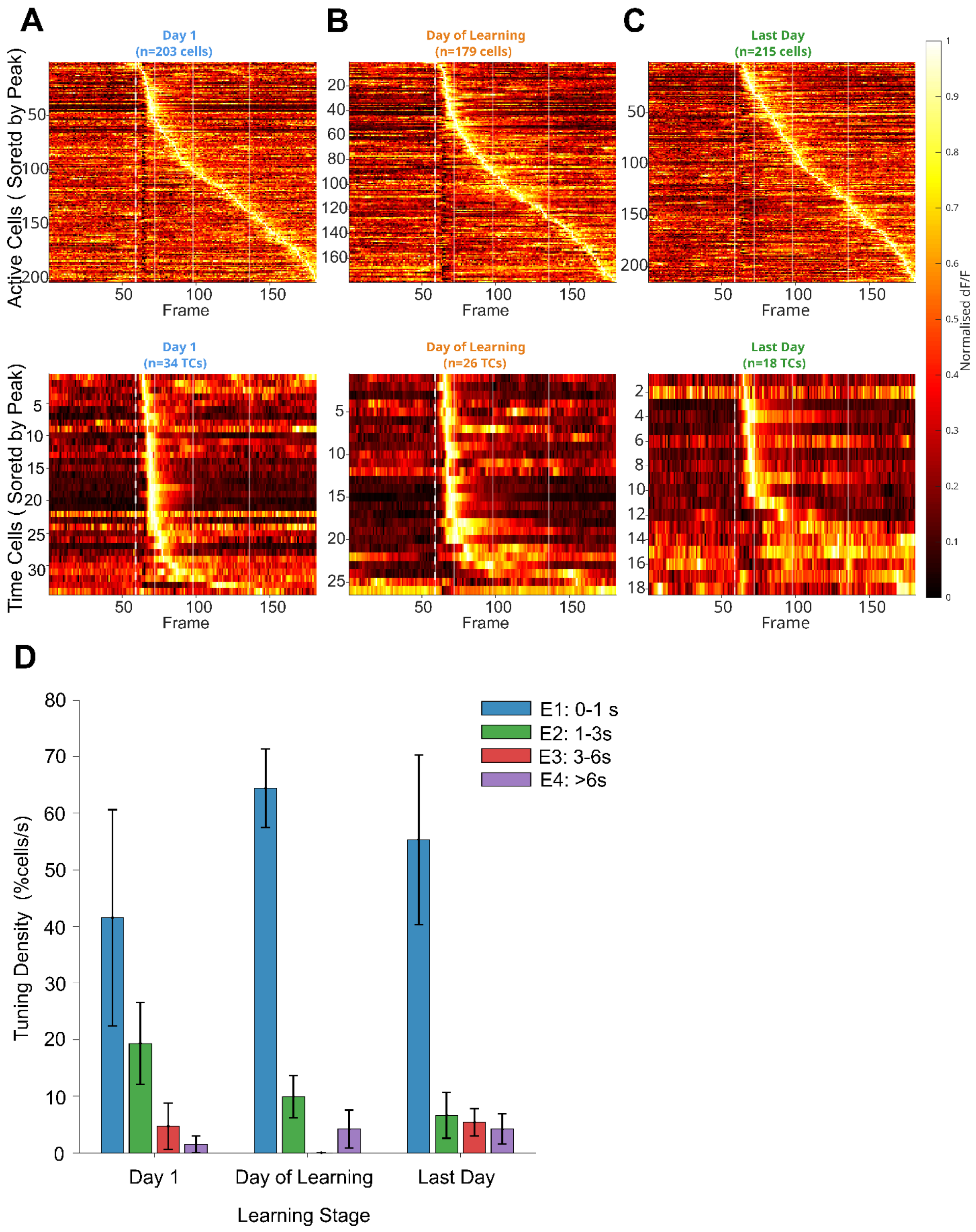

A
B
C
Day 1
(n=203 cells)
Day of Learning
(n=179 cells)
Last Day
(n=215 cells)
Active Cells ( Soretd by Peak)
Frame
Normalised dF/F
Day 1
(n=34 TCs)
Day of Learning
(n=26 TCs)
Last Day
(n=18 TCs)
Time Cells ( Soretd by Peak)
D
E1: 0-1 s
E2: 1-3s
E3: 3-6s
E4: >6s
Tuning Density (%cells/s)
Day 1
Day of Learning
Last Day
Learning Stage
